# Machine learning identifies cell-free DNA 5-hydroxymethylation biomarkers that detect occult colorectal cancer in PLCO Screening Trial subjects

**DOI:** 10.1101/2024.02.25.581955

**Authors:** Diana C. West-Szymanski, Zhou Zhang, Xiao-Long Cui, Krissana Kowitwanich, Lu Gao, Zifeng Deng, Urszula Dougherty, Craig Williams, Shannon Merkle, Matthew Moore, Chuan He, Marc Bissonnette, Wei Zhang

**Author notes:** Corresponding authors: Wei Zhang and Marc Bissonnette. Co-first authors: Diana West-Szymanski and Zhou Zhang.

## Abstract

**Background:** Colorectal cancer (CRC) is a leading cause of cancer-related mortality, and CRC detection through screening improves survival rates. A promising avenue to improve patient screening compliance is the development of minimally-invasive liquid biopsy assays that target CRC biomarkers on circulating cell-free DNA (cfDNA) in peripheral plasma. In this report, we identify cfDNA biomarker candidate genes bearing the epigenetic mark 5-hydroxymethylcytosine (5hmC) that diagnose occult CRC up to 36 months prior to clinical diagnosis using the Prostate, Lung, Colorectal and Ovarian (PLCO) Cancer Screening Trial samples.

**Methods:** Archived PLCO Trial plasma samples containing cfDNA were obtained from the National Cancer Institute (NCI) biorepositories. Study subjects included those who were diagnosed with CRC within 36 months of blood collection (i.e., case, n = 201) and those who were not diagnosed with any cancer during an average of 16.3 years of follow-up (i.e., controls, n = 402). Following the extraction of 3 - 8 ng cfDNA from less than 300 microliters plasma, we employed the sensitive 5hmC-Seal chemical labeling approach, followed by next-generation sequencing (NGS). We then conducted association studies and machine-learning modeling to analyze the genome-wide 5hmC profiles within training and validation groups that were randomly selected at a 2:1 ratio.

**Results:** Despite the technical challenges associated with the PLCO samples (e.g., limited plasma volumes, low cfDNA amounts, and long archival times), robust genome-wide 5hmC profiles were successfully obtained from these samples. Association analyses using the Cox proportional hazards models suggested several epigenetic pathways relevant to CRC development distinguishing cases from controls. A weighted Cox model, comprised of 32-associated gene bodies, showed predictive detection value for CRC as early as 24-36 months prior to overt tumor presentation, and a trend for increased predictive power was observed for blood samples collected closer to CRC diagnosis. Notably, the 5hmC-based predictive model showed comparable performance regardless of sex and self-reported race/ethnicity, and significantly outperformed risk factors such as age and obesity according to BMI (body mass index). Additionally, further improvement of predictive performance was achieved by combining the 5hmC-based model and risk factors for CRC.

**Conclusions:** An assay of 5hmC epigenetic signals on cfDNA revealed candidate biomarkers with the potential to predict CRC occurrence despite the absence of clinical symptoms or the availability of effective predictors. Developing a minimally-invasive clinical assay that detects 5hmC-modified biomarkers holds promise for improving early CRC detection and ultimately patient survival through higher compliance screening and earlier intervention. Future investigation to expand this strategy to prospectively collected samples is warranted.

## 1. INTRODUCTION

Colorectal cancer (CRC) is a leading cause of cancer-related mortality.^1^ According to the World Health Organization, more than 1.9 million new cases of CRC and more than 930,000 CRC deaths were estimated to have occurred globally in 2020.^2^ In the US, the 5-year relative survival is 65% for all stages combined, whereas the 5-year survival dramatically increases to 91% when the cancer is localized.^3^ Indeed, early detection and screening is associated with improved survival^4^ and many countries have implemented screening programs using fecal immunohistochemical tests (FITs), colonoscopies, or flexible sigmoidoscopies.^5, 6^

Although different screening methods have been recommended by U. S. Preventive Services Task Force,^6^ challenges of implementing an effective CRC screening program exist, including low patient adherence^7^, the invasiveness of methods like colonoscopy,^8^ and the inconvenience of procedures requiring dietary restrictions or multiple rounds of testing.^6^ While age, race, lifestyle factors (e.g., obesity, smoking history), diabetes, and a family history of CRC or certain genetic predispositions (e.g., Lynch syndrome, familial adenomatous polyposis syndrome) may provide an estimation for an individual’s risk for developing CRC, they lack sensitivity and specificity to be robust predictors.^9, 10^ Moreover, when individuals with average risk do not show any CRC symptoms, blood tests such as serum carcinoembryonic antigen (CEA) are not useful for screening.^11^ Therefore, the urgent development of convenient, minimally invasive, and potentially point-of-care screening assays remains a critical need.

Epigenetic modifications, specifically 5-hydroxymethylcystones (5hmC) have emerged as novel cancer biomarkers, given their relevance in regulating both tissue-specific gene expression^12^ and aberrant expression in cancers.^13^ Previous studies suggested that global 5hmC is differentially modified in various solid tumors, including CRC,^14–16^ indicating their potential for detection and as diagnostic biomarkers for CRC. Of particular interest is the development of a minimally-invasive blood-based test for CRC occurrence that targets differential epigenetic modifications associated with CRC development.^13^ Notably, our team has published a series of biomarker discovery studies^17^ in which 5hmC profiles in circulating cell-free DNA (cfDNA) derived from plasma demonstrated diagnostic values for clinically confirmed cancers such as CRC,^18^ stomach cancer,^18^ liver cancer,^19^ lung cancer,^20^ and glioma.^21^ However, it is not clear whether 5hmC in cfDNA can be exploited to detect CRC well before any clinical symptoms are manifested (i.e., pre-clinical or occult CRC “cases”).

In the current study, we collaborated with the National Cancer Institute (NCI) to obtain plasma samples from the Prostate, Lung, Colorectal and Ovarian (PLCO)^22^ Trial. The PLCO Trial was a randomized controlled trial to determine if cancer screening tests reduced cancer mortality. It enrolled ∼150,000 cancer-free participants across the United States from 1993 to 2001, collected 2.9 million biospecimens, and maintained systematic follow-up for 15 – 20 years.^22 23, 24^ We utilized this valuable biorepository to obtain archived plasma samples and identify 5hmC-modified biomarkers associated with CRC occurrence within three years after the sample collection. The PLCO samples provide a unique opportunity to explore early detection or develop a predictive model for CRC when clinical diagnosis is not suspected. Interestingly, previous studies have indicated the utility of protein biomarkers from PLCO samples for early detection of common cancers (e.g., CA19-9 for pancreatic cancer),^25^ thus supporting the utility of this precious resource for bridging the gap between early screening and current biomarker discovery efforts that are focused on biomarkers associated with “clinical cases.” In addition, by targeting novel genome-wide epigenetic signals associated with pre-clinical CRC cases, we expect to enhance our understanding of epigenetic pathways relevant to early development of CRC.

## 2. MATERIALS AND METHODS

### PLCO samples

In coordination with the NCI CDAS and Information Management Systems (supports the PLCO Trial dataset), a set of 603 PLCO Trial^23^ study participants (**Table 1**) that met our inclusion/exclusion criteria and 1:2 case:control design. “Cases” were defined as PLCO participants who developed CRC within 36 months from blood collection, i.e., early CRC or high-risk individuals with no symptoms and not diagnosed with CRC yet (n = 201). “Controls” were defined as those PLCO participants were not diagnosed with any cancer up until the patient was lost to follow-up after exiting the study (n = 402). And on average, the controls were followed for 16.3 years. All cases and controls were selected from the screening intervention arm of the PLCO Trial. Each “case set”, set of 2 controls and 1 case, were matched on age, sex, race/ethnicity, and time of sample collection. We excluded participants with non-CRC cancers, ulcerative colitis, Crohn’s disease, or Familial Polyposis. Most of the identified participants were of European ancestry (EA: European American) (173 cases and 346 controls), while the remainder were of African ancestry (AA: African American) (17 cases and 34 controls) or other race/ethnicity (11 cases and 22 controls). For quality control (QC) purposes, our study incorporated 40 control sample replicates, consisting of 27 within-batch and 13 across-batch replicate samples.

**Table 1.** Clinical and demographical information of the PLCO samples.

|  | **Cases** | **Controls** | **Total** |
| --- | --- | --- | --- |
| No. | 201 | 402 | 603 |
| Age (Years) (mean ± std) | 64.2 ± 4.8 | 63.9 ± 5.1 | 64.0 ± 5.0 |
| Sex |  |  |  |
| Female | 93 | 185 | 278 |
| Male | 108 | 217 | 325 |
| Population |  |  |  |
| EA | 173 | 346 | 519 |
| AA | 17 | 34 | 51 |
| Others | 11 | 22 | 33 |
| BMI (mean ± std) | 27.9 ± 5.0 | 27.7 ± 4.7 | 27.8 ± 4.8 |
| Smoking history |  |  |  |
| Current | 22 | 33 | 55 |
| Never | 94 | 193 | 287 |
| Former | 85 | 176 | 261 |
| CRC Stage |  |  |  |
| Stage I/II | 110 | - | 110 |
| Stage III/IV | 87 | - | 87 |
| NA | 4 | - | 4 |
| Time to diagnosis: (mean±std) Years | 1.1 ± 0.9 | - | - |
| Average follow-up (mean±std) Years | 12.3 ± 5.0 | 16.3 ± 3.1 | 15.3 ± 4.1 |

Notes: CRC, colorectal cancer; EA, European American; AA, African American; BMI, body mass index; std, standard deviation. Other populations include Asians and Latinos.

Under a cooperative agreement, NIH CDAS provided de-identified information, including CRC diagnosis and stage, self-reported race/ethnicity (i.e., population), age, time between blood collection and CRC diagnosis (≤36 months), sex, and other relevant variables (e.g., BMI [body mass index], smoking history). CRC diagnosis and staging was confirmed at individual screening centers according to the American Joint Committee on Cancer (AJCC) guidelines (5^th^ edition at the time of the trial, and 7^th^ edition calculated by CDAS post-trial). The samples were aliquoted and organized in assay batches that included an inter- and intra-batch technical replicates, then shipped to The University of Chicago. The PLCO Trial obtained informed consent from each participant.^23^ This study was approved by the Institutional Review Board of The University of Chicago.

### Genome-wide 5hmC profiling

We obtained frozen aliquots of EDTA plasma samples (200 – 300 µL per study participant) from the PLCO repository. Samples were immediately stored at -80°C until processing. CfDNA was extracted from the plasma samples using the QIAamp Circulating Nucleic Acid Kit (Qiagen) and approximately 3 – 8 ng of cfDNA was isolated from each plasma sample for use in the 5hmC-Seal assay. The 5hmC-Seal protocol^18, 26^ was then used to selectively label and enrich 5hmC-contatining cfDNA fragments. Briefly, T4-beta-glucosyltransferase (Thermo) was used to modify the 5hmC modifications on cfDNA with UDP-azide-glucose (Active Motif). Next, we performed a click reaction between the azide-linked modifications (products from the previous reaction) and the strained alkyne DBCO-PEG_4_-biotin (Click Chemistry Tools). The resulting biotinylated-5hmC-modified cfDNA fragments were then affinity captured using Dynabeads M-270 Streptavidin (Thermo), and libraries were constructed using on-bead PCR amplification using the KAPA HyperPlus kit (Roche). The constructed libraries were then sequenced using the Illumina NovaSeq 6000 platform (paired-end 50 bp reads) at The University of Chicago Functional Genomics Core.

### Bioinformatic processing

The basic bioinformatic processing aimed to provide a robust summarization of the 5hmC-Seal data as well as facilitate data analysis and result interpretation, as described in our previous publications.^17–19^ Adapter sequences were removed from raw sequencing reads using Trimmomatic.^27^ Low quality bases at the 5’ and 3’ were trimmed based on phred score to a minimum length of 30 bp. The sequencing reads were then aligned to the human genome reference (hg19) using Bowtie2 with the end-to-end alignment mode.^28^ Read pairs were concordantly aligned with fragment length ≤ 500 bp and with average ≤ 1 ambiguous base and up to four mismatched bases per 100 bp length. Alignments with Mapping Quality Score ≥ 10 were counted based on the GENCODE^29^ gene body annotations using featureCounts,^30^ without strand information. The raw read counts were then normalized using DESeq2,^31^ which performs an internal normalization that corrects for library size.

### Quality control testing

Our study incorporated 40 QC sample replicates, consisting of 27 within-batch and 13 across-batch QC samples. For additional replicates to test the technical robustness of the assay in these limited cfDNA PLCO samples, we compared profiling data with our previously-sequenced 5hmC dataset (n = 27 pre-clinical cases and n = 28 controls) from a separate project.^32^

### Cellular deconvolution

Cellular deconvolution on the summarized genome-wide 5hmC profiles was performed to estimate relative contribution of a variety of tissue types and white blood cells using the Cibersortx tool,^33^ which determines relative cell type abundance from bulk tissues with digital cytometry. Specifically, we assembled a list of up- and down-modified genes for 19 tissues from our previously published 5hmC Tissue Map^12^ and 8 white blood cell types from Nakauchi et al.^34^ as the reference input for Cibersortx. The estimated proportions that are scaled to a sum of 100% were reported to represent the relative contribution of a particular tissue/cell type to the genome-wide 5hmC profiles generated from each cfDNA sample.

### Detection of 5hmC signatures associated with pre-clinical CRC

The PLCO samples were randomly split at 2:1 ratio into a training set (134 cases and 269 controls) and a validation set (67 cases and 133 controls). To enhance downstream modeling efficiency and reduce overfitting, we first used multivariable Cox proportional hazards models adjusting for age, sex, race/ethnicity, BMI, and smoking status, to detect a list of most informative 5hmC signatures (gene bodies) associated with pre-clinical CRC cases in the training samples (p < 0.05) and showed a trend for the same direction of association in the validation samples. The resulting candidate list of pre-clinical CRC-associated 5hmC marker genes were investigated for functional enrichment of Gene Ontology biological processes and canonical pathways maintained by the KEGG (Kyoto Encyclopedia of Genes and Genomes) database,^35^ using clusterProfiler 4.0 tool.^36^

### Machine learning modeling of a 5hmC-based algorithm for predicting CRC occurrence

We applied a two-step procedure in the training samples to develop a weighted predictive score (wp-score) for distinguishing patients who developed CRC from non-cancer controls. In Step 1, as described in the previous section, the multivariable Cox models adjusting for age, sex, race/ethnicity, BMI, and smoking status, were used to select a list of candidate 5hmC marker genes associated with pre-clinical CRC cases (or those features more likely to be informative). In Step 2, feature selection was performed using the elastic net regularization that aimed to fit a multivariable Cox model for predicting CRC occurrences over time using the *glmnet* library.^37^ The elastic net model was cross-validated (10-fold) with α value of 0.5, indicating a mix of LASSO and ridge regression penalties. The optimal λ was determined that minimized the cross-validated error. The coefficients of the model, estimated at the optimal λ, were extracted. These coefficients represent the strength and direction of the association between each feature and the survival outcome, under the optimal level of regularization. Only features with non-zero coefficients were retained. This selection process was repeated 100 times by bootstrap, and the gene bodies selected in at least 95% of the iterations were retained to fit the final Cox model. The final Cox model was applied to the 5hmC levels of selected gene bodies, and a wp-score was calculated as: wp-score = 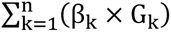 where for the *k*th gene, *β_k_* is the final Cox model coefficient and *G_k_* is the normalized 5hmC levels. The area under the curve (AUC) and 95% confidence intervals (CI) were computed (e.g., for periods <6 months, 6-12 months, 12-24 months, and 24-36 months between CRC diagnosis and blood collection) to show model performance.^38^ The performance of a single model component was evaluated using AUC. Sensitivity and specificity were computed using the wp-score at different cutoffs that achieved a range of specificities in the training samples. We also evaluated predictive performance by combining the 5hmC-based wp-scores and other risk factors such as age, sex, BMI, and smoking status.

In addition, using RNA-seq data from The Cancer Genome Atlas (TCGA) Project (CRC tumors, n=469; normal tissues, n=41), we investigated whether the final model components based on 5hmC exhibited gene dysregulation in CRC tumors.^39^

### Prediction of CRC risk over time with the wp-scores

A clinical event was defined as the diagnosis of CRC. Individual cfDNA samples were grouped into two risk groups based on the median of wp-scores in the training and validation samples: high-risk individuals comprised of those individuals with higher risk (greater than the median) for future CRC occurrence and low-risk individuals comprised of those individuals less likely to develop CRC in the future. Kaplan-Meier (KM) analysis was performed, and hazard ratios (HR) were calculated based on the final multivariable Cox models.

### Statistical analysis

All statistical analyses were performed using the R Statistical Computing Environment (v4.1.2).^40^ When testing the relationship between wp-scores and clinical variables, continuous variables were analyzed using the two-sided Student’s *t*-tests, and categorical variables were analyzed using the Chi-square test. The *limma* R package^41^ was used to detect differentially expressed model genes between TCGA CRC tumors and normal tissues, and a false discover rate (FDR) < 0.05 was considered statistically significant. T-SNE was used to visualize the TCGA datasets based on the differential expressed genes. To assess the discriminative performance of the wp-scores, a receiver operating characteristic (ROC) curve was generated, and an AUC was computed with 95% confidence intervals (CIs) using ROCR R package.^38^ Statistical significance for the KM analysis was determined using the log-rank test.

## 3. RESULTS

### 5hmC profiling of PLCO samples

In collaboration with the National Cancer Institute (NCI) Cancer Data Access System (CDAS) we designed a study using PLCO samples in which we defined pre-clinical “cases” as participants with bioarchived plasma collected within 3 years of their CRC diagnosis (n = 201), and non-cancer “controls” as those who remained cancer-free until they were lost to follow-up after exiting the study (n = 402). Cases and controls were matched by age, race, and gender. **Table 1** shows the demographics and clinicopathologic characteristics of the cases and controls, and **Figure 1** shows the flow chart of sample selection, assaying, and biomarker identification. Additionally, we requested replicate samples for within- and across-batch quality control (n = 40).

**Figure 1.**
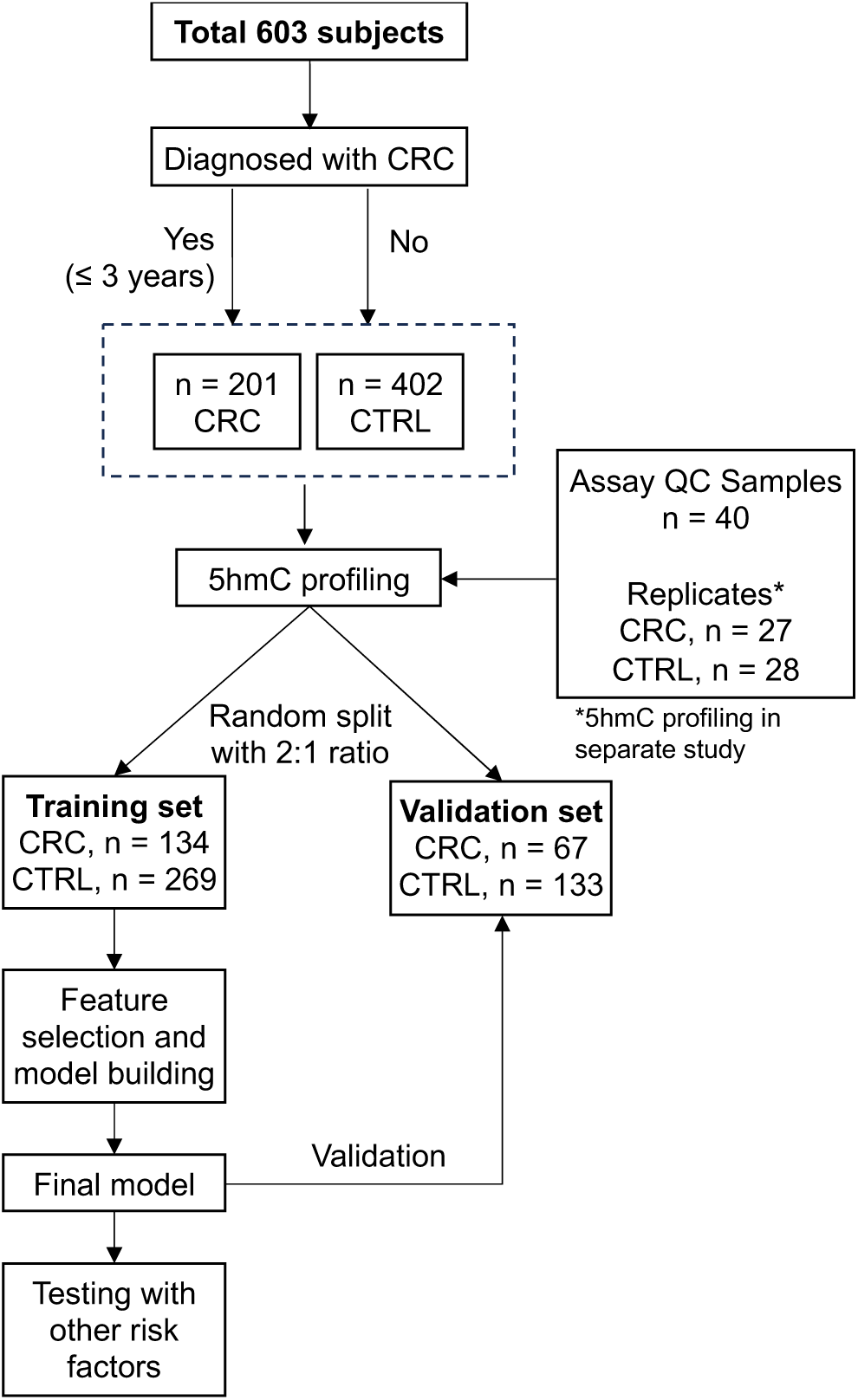
Study design and workflow. For the current project, we obtained 603 cfDNA samples from the PLCO repository, including pre-clinical CRC cases and age-, sex-, race/ethnicity (population)-matched controls. Genome-wide 5hmC profiles were obtained using the 5hmC-Seal technique and the next-generation sequencing (NGS), followed by association analysis and statistical modeling. In addition, QC samples were used to test between- and within-batch variability, and replicate CRC and CTRL samples were used for assay robustness. CRC: colorectal cancer pre-diagnostic (Dx) cases; CTRL: non-cancer controls; cfDNA: cell-free DNA; 5hmC: 5-hydroxymethylcytosine; PLCO: The Prostate, Lung, Colorectal and Ovarian Cancer Screening Trial.

PLCO samples were aliquoted, randomized by case set, and organized into study batches before shipment to The University of Chicago. Technicians were blinded to the case or control status of the samples during the cfDNA extraction and 5hmC-Seal assay. Following next-generation sequencing, ∼19 million reads were obtained for each PLCO sample from both cases and controls (**Figure S1A**). The principal components analysis (PCA) of the genome-wide 5hmC data indicated no apparent outliers across all cfDNA samples (**Figure S1B**), indicating no systematic bias due for example to different experimental batches. The PLCO samples were randomly split at 2:1 ratio into a training set (134 cases and 269 controls) and a validation set (67 cases and 133 controls). Age, sex-, and race/ethnicity distributions shown in **Figure S2** suggested no systematic biases regarding these variables between training and validation samples: age (64.0±5.0 vs. 63.8±4.9 years), sex (53.8% vs. 54.0% males), and race/ethnicity (85.6% vs. 87.0% EA).

### Cellular deconvolution of 5hmC profiles in cfDNA

Taking advantage of 5hmC cell/tissue-type specific reference databases developed by our team^12^ and Nakauchi et al,^34^ we performed cellular deconvolution on the summarized 5hmC data to estimate relative contributions of various tissues and white blood cells. The estimated abundance (i.e., proportion) was reported to represent the relative contribution of a particular tissue or cell type in cfDNA for each sample. Of note, there was not a statistically significant trend of increased or decreased colon-derived signals when the blood collection was closer to CRC diagnosis (e.g., CRC diagnosed within 6 months vs. CRC diagnosed after 24 months) (**Figure S3**), suggesting that at the time of blood collection, early-stage colon cancer was “silently” developing, with limited colon-derived tissue cfDNA released into the circulation. Overall, the failure of cellular deconvolution to suggest occult CRC development (via colon-derived tissue cfDNA) underscores the need to investigate gene-specific signals that differentiate cases from controls in these pre-clinical samples.

### Detection of 5hmC signatures associated with pre-clinical CRC

Multivariable Cox proportional hazards models identified 890 gene bodies with 5hmC modification levels significantly associated with pre-clinical CRC cases in the training set (p < 0.05). Examining the validation set, 540 gene bodies were further confirmed in association with pre-clinical CRC cases (**Figure 2A**; **Table S1).** Functional annotation of these 540 pre-clinical CRC-associated genes suggested enrichment of multiple KEGG signaling pathways, including calcium and Rap1 signaling pathways, and cancer related pathways (**Figure 2B**; **Table S2**).

**Figure 2.**
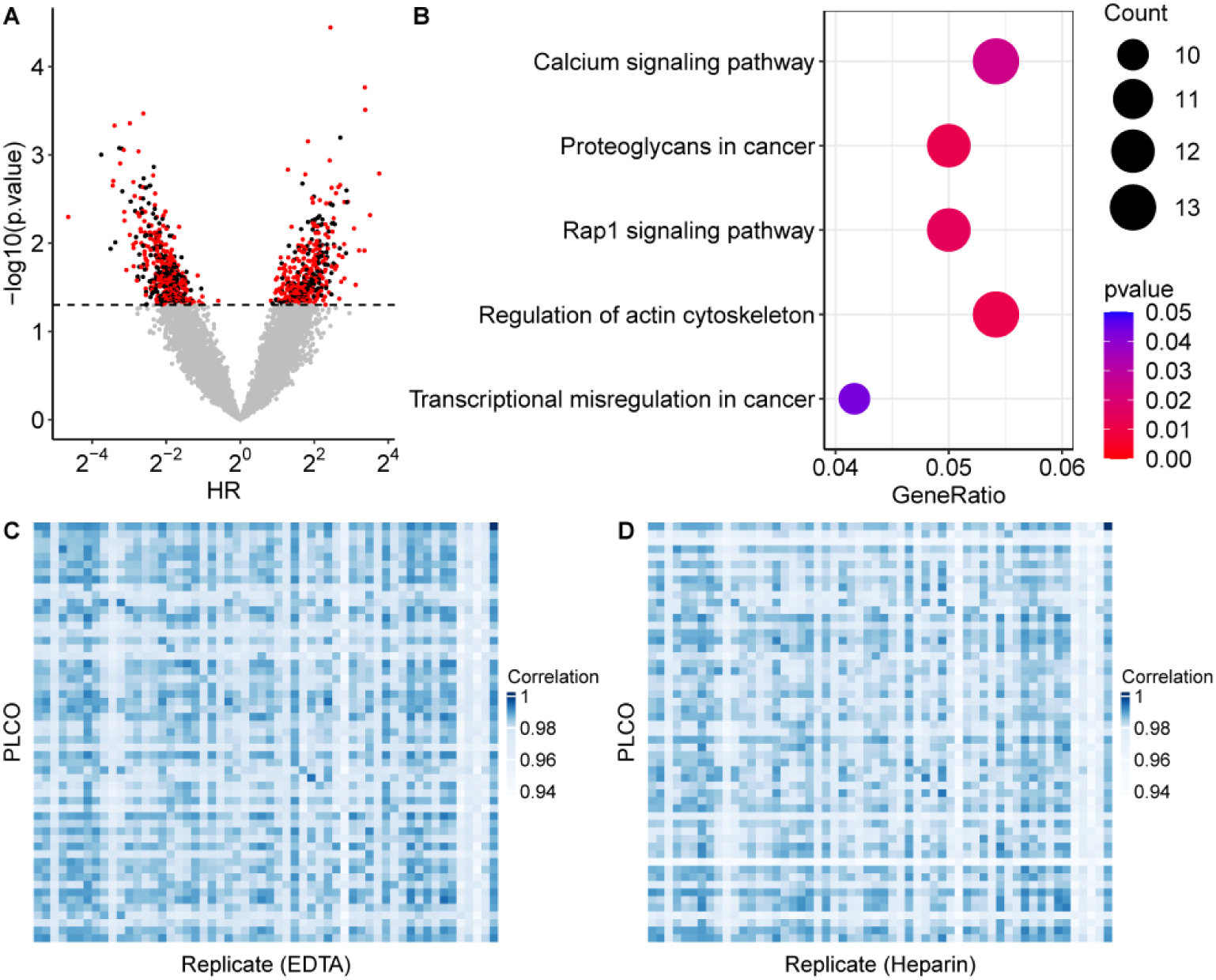
Identification of 5hmC signatures associated with pre-clinical CRC cases. The univariate Cox proportional hazards model is performed on each 5hmC marker gene (i.e., gene body) to identify differential signatures associated with pre-clinical CRC cases. **A.** The volcano plot shows hazard ratios (HR) in the X-axis and p-values in the Y-axis in the training samples. Those gene bodies with the same association direction in the validation samples are highlighted in red. **B.** The KEGG pathways enriched in the 5hmC signatures associated with pre-clinical CRC cases. KEGG: Kyoto Encyclopedia of Genes and Genomes. **C-D.** A correlation matrix shows technical robustness of the 5hmC-Seal assay between PLCO samples and duplicates stored in: EDTA (**C**) and Heparin (**D**).

The QC samples demonstrated a higher correlation with their corresponding replicates (mean Pearson’s r = 0.9799) compared to other samples (mean r = 0.9758), achieving statistical significance with a p-value of 0.036. Additionally, a subset of this study’s subjects (n = 27 cases and n = 28 controls) overlapped with study subjects for a previous technical development study^32^ in which we tested EDTA and heparin plasma anti-coagulants. When comparing genome-wide 5hmC distributions between our current sequencing data to the previous sequencing dataset to genome-wide 5hmC distributions in our current study, we found a significantly higher correlation between the same PLCO participants compared to random pairs of samples stored in EDTA (**Figure 2C**) or Heparin (**Figure 2D**), demonstrating technical robustness of the 5hmC-Seal assay.^32^

### A 5hmC-based model for predicting risk for CRC in pre-clinical samples

For those CRC cases, 55.0% were diagnosed with stages I-II CRC in the training samples, compared to 57.6% in the validation set (**Figure S2D**). Using machine learning, a 32-gene model was developed in the training samples to predict CRC occurrence (i.e., those individuals diagnosed with CRC within 36 months of blood collection) (**Figure 3A, Table S3**). Several model genes, such as *TGFB2, TLN2*, and *KMT2A* were found to be involved in enriched KEGG pathways (e.g., *TGFB2* in the “proteoglycans in cancer pathway”, and *TLN2* in the “Rap1 signaling pathway”, and *KMT2A* in the “transcriptional misregulation in cancer pathway”) (**Table S2**). The weighted prediction scores (wp-scores) computed based on this 32-gene model showed high capacity for distinguishing cases and controls in both training (AUC: 77.0%, 95% CI: 72.1-81.8%) and validation (AUC: 72.8%, 95% CI: 65.8-79.7%) samples (**Figure 3B**). Using the wp-score cutoff (i.e., -43.48) that maximized the Youden’s index in the training set, we achieved 0.58 sensitivity and 0.83 specificity in the training samples, compared to 0.47 sensitivity and 0.73 specificity in the validation samples. Examination of the predictive value of the 5hmC-based wp-scores for CRC occurrence showed comparable model performance regardless of race/ethnicity (population), tumor stage at diagnosis, age, sex, obesity as defined by BMI, and smoking history (**Figure 3C-H**).

**Figure 3.**
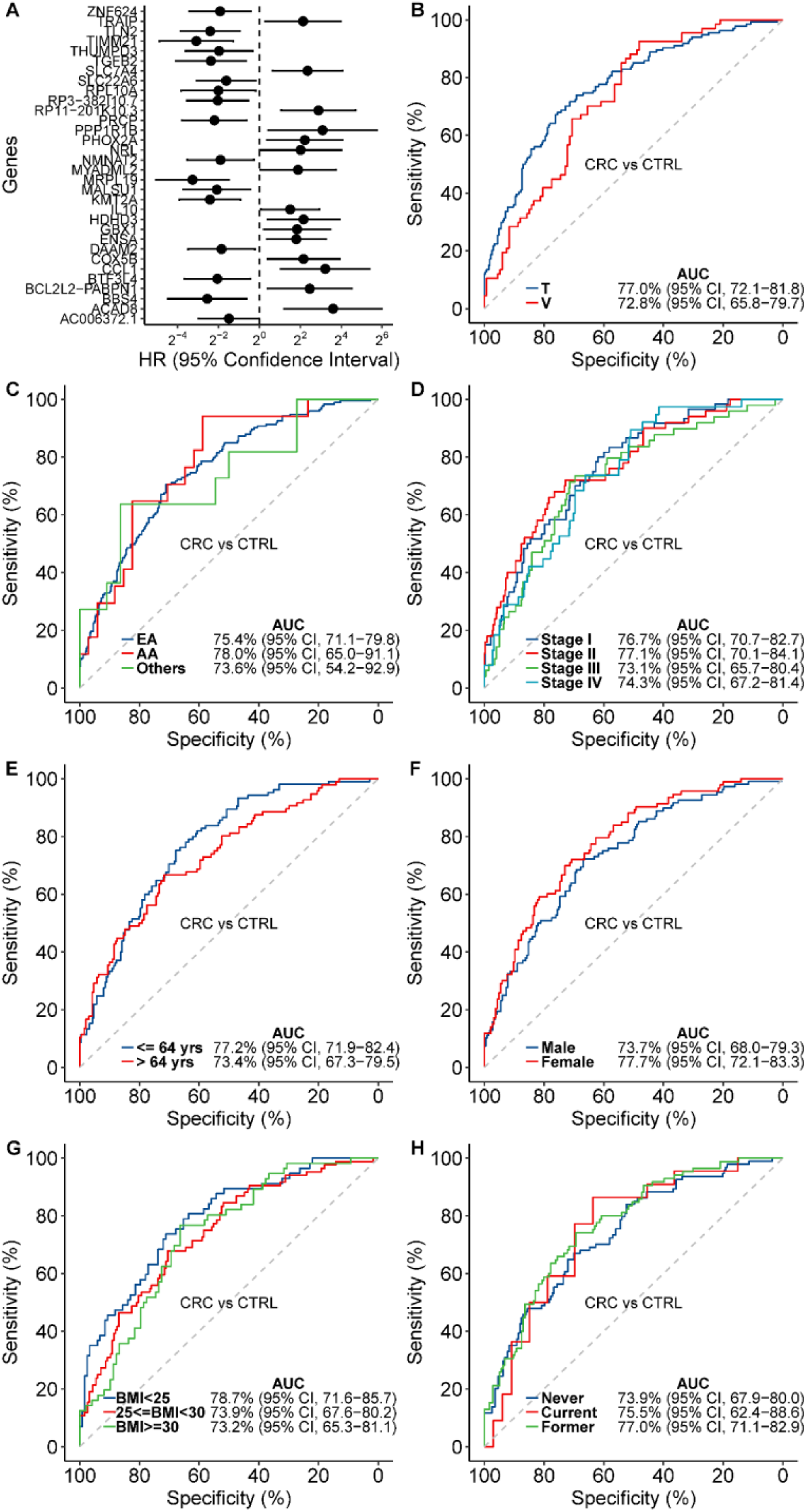
Performance of a 5hmC-based model for predicting risk for CRC in pre-clinical samples. The elastic net regularization, a machine learning approach was used to perform feature selection under the Cox proportional hazard model. **A.** A predictive model for CRC occurrence in pre-clinical samples comprised of a panel of 17 genes. Shown is a forest plot for the model components and individual hazard ratios (HR). **B.** Shown is the performance of the 5hmC-based predictive model for CRC occurrence in pre-clinical samples in the training (T) and validation (V) samples. AUC: area under the curve; CI: confidence interval. **C-H** show the performance of the 5hmC-based predictive model for CRC occurrence by various risk factors, including: **C.** race/ethnicity (population); **D.** CRC stage; **E.** age; **F.** sex; **G.** obesity as defined by BMI; and **H.** smoking history. BMI: body mass index.

Of note, although these common risk factors (e.g., age, lifestyle factors) did not show predictive value for CRC occurrence in the pre-clinical PLCO samples, a general trend of increased model accuracy in terms of AUC for wp-scores was achieved when CRC diagnosis was closer to blood collection in all samples (**Figure 4A**). Conversely, age, BMI, and smoking history did not exhibit this AUC trend as the time to diagnosis were closer (**Figure 4B-C**). Together, these data suggest that the machine learning-derived panel of 32 differentially 5hmC-modified genes provides a better predictor of CRC than other known CRC risk factors.

**Figure 4.**
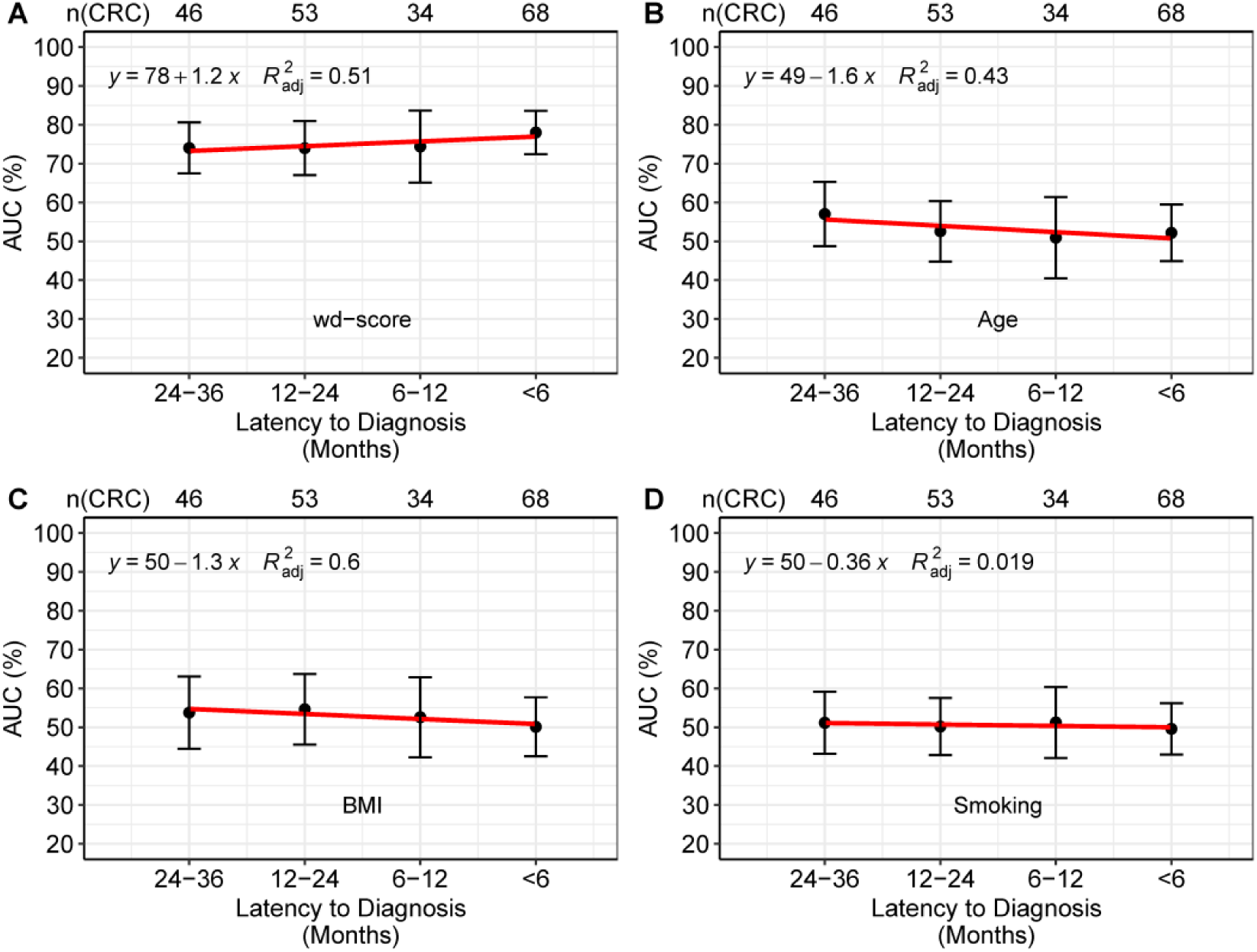
comparison of predictive value between wp-scores and risk factors for CRC. Predictive performance is shown for the 5hmC-based wp-scores and various risk factors for CRC, by different time periods between blood collection and CRC diagnosis: ≤ 6 months, 6-12 months, 12-24 months, and 24-36 months. For each time period, 95% CI is also shown for AUC. **A.** wp-scores; **B.** age; **C.** obesity; and **D.** smoking history. AUC: area under the curve; CI: confidence interval.

### Risk stratification with the 5hmC-based wp-scores for CRC occurrence in pre-clinical samples

We next sought to determine if the 5hmC-based wp-scores showed predictive value for CRC risk (event-free survival) over time by the Kaplan-Meier (KM) analysis. Specifically, high-risk individuals (i.e., those with wp-scores higher than the median) showed a significantly higher risk for CRC occurrence than low-risk individuals (i.e., those with wd-scores lower than the median) in the training (HR: 3.3, 95% CI: 2.6-5.8, log-rank test p-value < 0.0001) and validation (HR: 3.1, 95% CI: 1.8 - 5.8, log-rank test p-value < 0.0001) samples (**Figure 5A-B**). Comparing with other risk factors, such as smoking, sex, race, BMI, and age, the wp-score showed significant predictive value in terms of hazard ratios (HR) in training and validation samples (**Figure 5C-D**). We also assessed a complete model, which integrates the wp-score with the previously mentioned risk factors, for predicting CRC occurrence within the training and validation sets. However, this integration did not enhance the predictive capability (**Figure 5E**). A nomogram comprised of the 5hmC-based wp-scores as well as age, sex, obesity, race, smoking history was constructed to provide an overview of the impact of predictive value for individual predictors (**Figure 5F**), indicating the wp-score accounts for the majority of the predictive points, while other risk factors offering little to no additional contribution.

**Figure 5.**
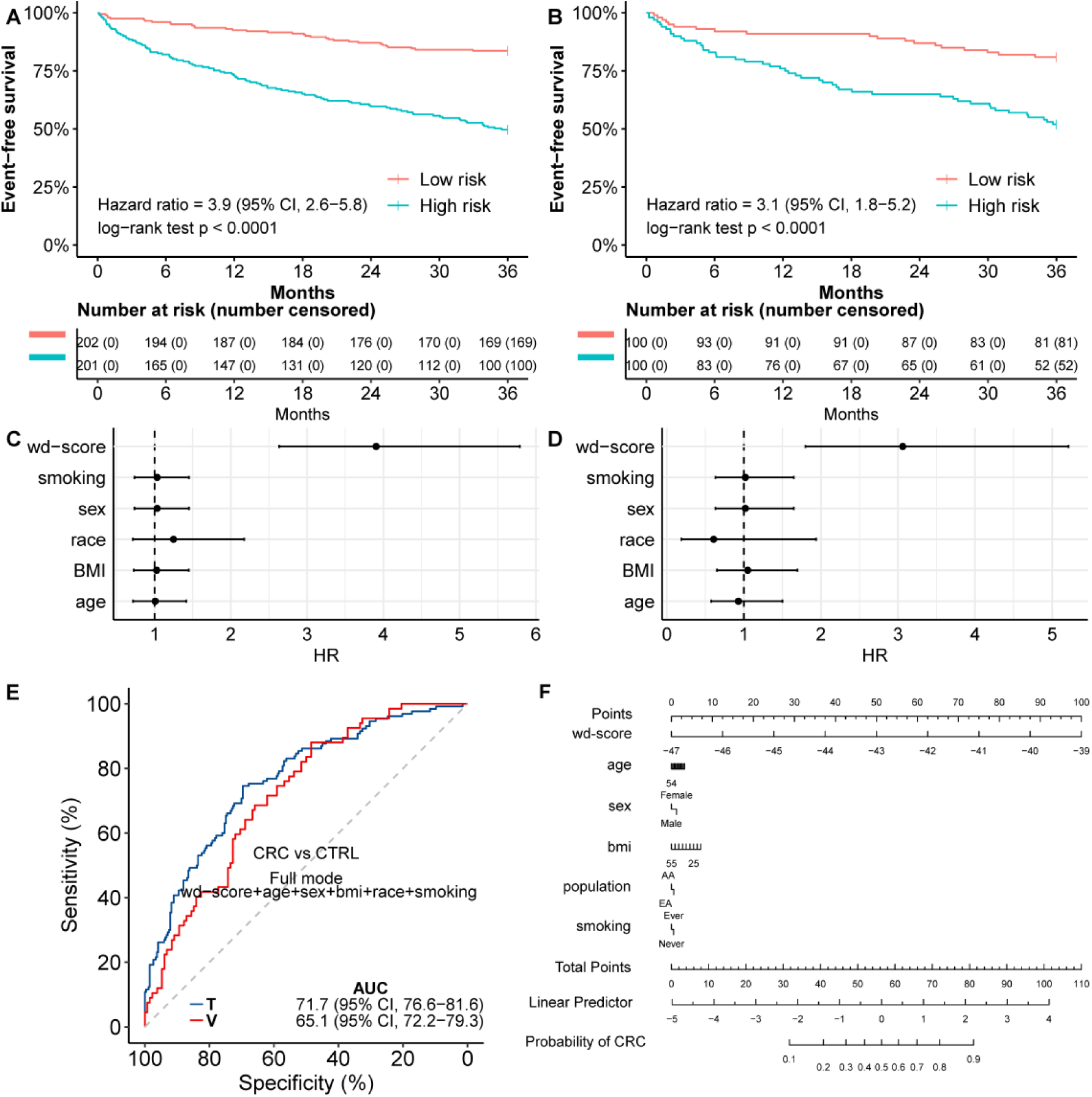
Risk stratification with the 5hmC-based wp-scores for CRC occurrence in pre-clinical samples. The wp-scores computed based on the final predictive model for CRC occurrence are used to assign individuals samples into a high-risk group (i.e., those with wp-scores higher than the median) and a low-risk group (i.e., those with wp-scores lower than the median). Kaplan-Meier (KM) plots show the differential rate of CRC occurrence over time (months) according to the 5hmC-based risk in **A.** training samples, and **B.** validation samples. Forrest plot showing the hazard ratios (HR) for wp-score and other risk factors in **C.** training samples; **D.** validation samples. **E.** Shown is the performance of full model integrating wp-score and other risk factors for predicting CRC occurrence in pre-clinical samples in the training (T) and validation (V) sets. **F.** A nomogram for risk evaluation. Relative contributions of the 5hmC-based wp-scores as well as age, sex, obesity, smoking history are shown.

### Model genes demonstrated relevance in gene expression based on data from TCGA

For 28 out of the 32 final model component genes, the normalized RNA-seq data of 469 primary tumors and 41 normal tissues were extracted from TCGA CRC dataset. Differential analysis showed that 10 (35.7%) out of these 28 model genes were differentially expressed between tumors and normal tissues (**Table S4**, **Figure S4**), indicating the relevance to tumor transcriptional dysregulation of these 5hmC markers in cfDNA in an independent set of TCGA samples. For example, higher expression of DAAM2 (Disheveled-associated activator of morphogenesis 2) has been indicating an unfavorable prognosis of CRC,^42^ And ACAD8 (Acyl-CoA Dehydrogenase Family Member 8) showed favorable prognosis in CRC according to Human Pathology Atlas.^43^

## 4. DISCUSSION

CRC screening has proven effective in improving clinical outcomes and reducing CRC-related mortality.^4^ The American Gastroenterological Association recommends that screening for individuals at average-risk for CRC should begin at age 45 years, with a screening colonoscopy every 10 years being the most effective screening modality.^44^ However, patient compliance rates remain suboptimal, partly due to complications associated with colonoscopy preparations.^45, 46^ Several minimally invasive approaches have been developed to complement screening colonoscopies that could potentially enhance patient compliance, especially those exploiting blood (e.g., *SEPT9* promoter methylation)^47^ or stool (e.g., FIT, Cologuard).^48, 49^ Compared to stool-based tests, studies suggest that blood-based early detection or screening tools might achieve an even higher level of patient compliance.^50, 51^ Although there have been significant advances in blood-based tests for early CRC detection, there are limitations including relatively low sensitivity (e.g., the *SEPT9* test) or a biomarker discovery strategy that targeted diagnosed CRC cases. In the current study, our primary goal was to take advantage of the unique resource from the PLCO Trial and our team’s highly sensitive 5hmC-Seal technique to identify 5hmC-modified biomarker candidates on circulating cfDNA in pre-clinical samples obtained from individuals for whom CRC diagnosis occurred months or even years after blood collection. To our knowledge, this study is the largest effort exploiting the PLCO resources to address the gap in CRC biomarker discovery that traditionally has been focused on biospecimen collected when patients were already diagnosed.

All of the CRC “cases” in our study were diagnosed up to 36 months after the blood was collected, during which no symptoms or abnormalities were detected in the clinic. As expected, cellular deconvolution revealed no significant bias in tissue origin between cases and controls, across a variety of tissue and cell types, including immune cells and colon tissues. Of note, the absence of significant changes in colon-derived signals near CRC diagnosis underscores the subtle progression nature of early-stage CRC, which remains undetectable due to minimal colon-derived cfDNA release into the circulation (e.g., CRC developed within 6 months vs. after 24 months). This observation highlights to the challenges associated with early CRC detection and the limitations inherent in cellular deconvolution techniques. This stresses the need to investigate gene-specific signals in pre-clinical PLCO samples for potential biomarker discovery, as demonstrated in our current study.

Our primary goal in this research was to enhance the early detection and risk stratification of CRC through a more comprehensive characterization of 5hmC profiles in cfDNA. The 32-gene model we developed, informed by specific 5hmC modifications, represents a significant advancement by predicting CRC occurrence up to 36 months prior to diagnosis. Our early prediction capability suggests an improvement over existing CRC screening methods which typically detect cancer closer to or at the time of clinical diagnosis, although head-to-head comparisons would be needed to confirm this. Our model’s performance across diverse populations, regardless of variations in BMI, smoking history, and race/ethnicity, further underscore its potential for widespread clinical implementation. Moreover, the associated gene bodies and pathways we identified, such as calcium and Rap1 signaling pathways as well as cancer-related pathways, provide deeper biological insights into early CRC pathogenesis. These results open new avenues for the development of more targeted, minimally-invasive screening strategies that might significantly improve early detection rates and, consequently, patient outcomes in CRC.

It is important to acknowledge several key areas that warrant further investigation and refinement. Firstly, the prolonged storage time of the PLCO samples, while invaluable for our current analysis, raises the need for validation in fresher samples to ascertain the stability and consistency of our findings. Secondly, our study was not equipped to thoroughly evaluate disparities across different populations, which is a critical aspect for ensuring the broad applicability of our model. Additionally, the relatively modest sample size, while sufficient for our current analytical design, presents a balancing act between achieving robustness in analysis and avoiding overfitting. This limitation points to the necessity of expanding our study to include larger and more diverse cohorts. Future work should also consider incorporating heparin-stored resources, which could offer a broader scope and provide many additional PLCO samples for further validation of our findings. In this regard, we have recently shown that heparin storage is compatible with the 5hmC-Seal assay.^32^ Looking ahead, the potential for developing a predictive models for additional cancers based on our approach is both exciting and challenging, necessitating careful consideration of various genomic and epigenetic factors, including but not limited to epigenetic changes associated with aging. These limitations, while underscoring the need for caution in interpreting our current results, also pave the way for exciting future research directions that could significantly advance the field of cancer diagnostics and personalized medicine.

In conclusion, our study marks a significant step forward in the early detection and risk stratification of colorectal cancer through advanced analysis of 5hmC profiles in cfDNA. Our research demonstrates the potential of a 32-gene model for predicting CRC well in advance of traditional diagnostic methods. This work not only offers a promising avenue for minimally-invasive cancer screening, but also lays the groundwork for future studies aimed at refining and expanding the application of epigenetic profiling in cancer diagnostics.

## ACKNOWLEDGEMENTS

This work was partially supported by the NCI (U01CA217078). This work was partially supported by the Institute for Translational Medicine and the Chicago Biomedical Consortium. We would also like to thank the NCI for access to the data collected by the PLCO Trial that was funded in whole or in part with federal funds from the NCI, US National Institutes of Health (NIH). Chuan He is an investigator of the Howard Hughes Medical Institute.

