## Supplementary Figures and Tables for "Machine learning identifies cell-free DNA 5-hydroxymethylation biomarkers that detect occult colorectal cancer in PLCO Screening Trial subjects"

A

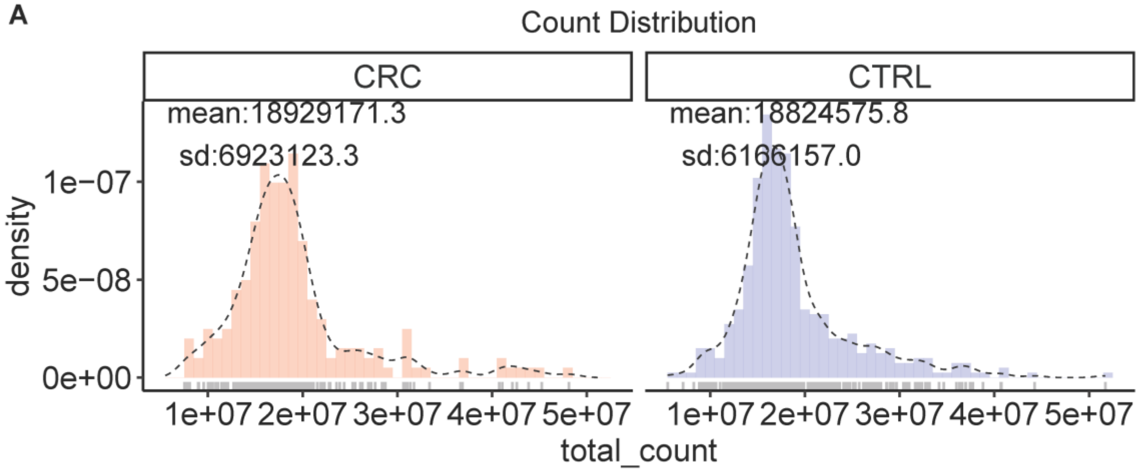


B

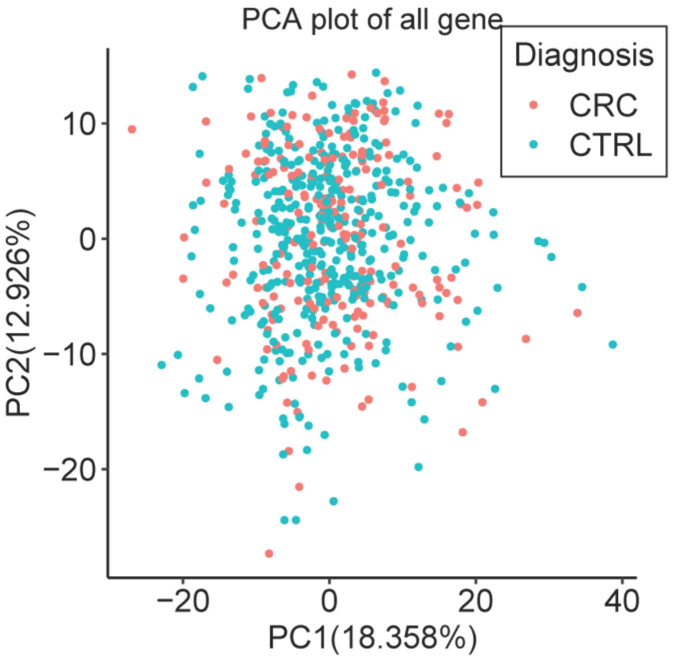


**Figure S1. Overview of the genome-wide 5hmC data generated from the PLCO samples.**

The genome-wide 5hmC profiles of 603 PLCO samples generated using the 5hmC-Seal and the next-generation sequencing (NGS) are shown to provide an overview of the profiling results. **A.** The read count distribution between cases (CRC) and controls (CTRL). Only unique reads are counted. **B.** The principal components analysis (PCA) plot of the PLCO samples using genome-wide 5hmC data. No apparent outliers exist in either cases (CRC) or controls (CTRL).

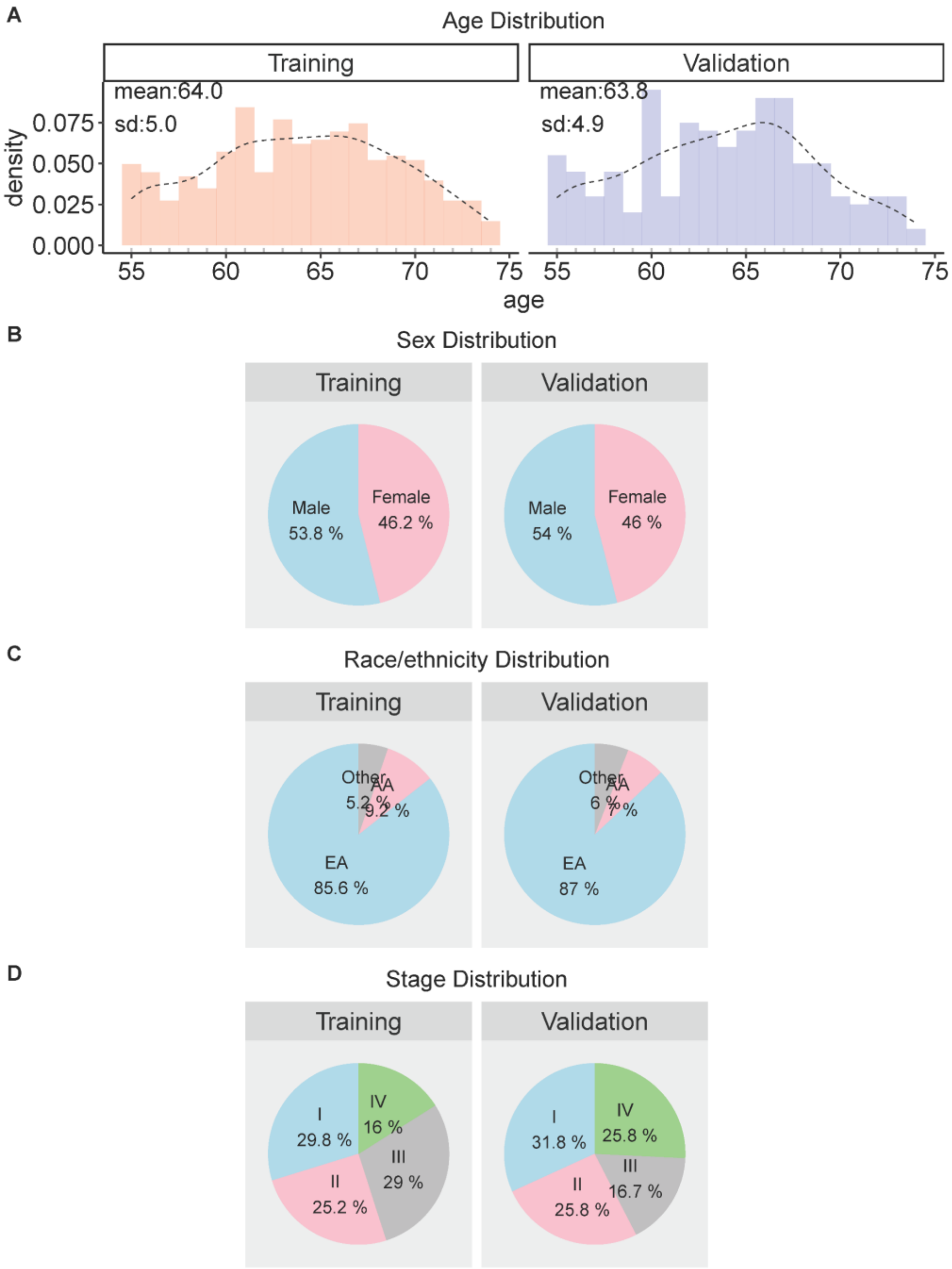


**Figure S2. Distributions of demographic variables between training and validation samples.**

The training set is comprised of 134 cases and 269 controls randomly selected from the final analytical set (66.7%) that excluded outliers and those that failed the 5hmC-Seal profiling. The remaining 67 cases and 133 controls (33.3%) are combined in the validation set. The distributions of the following demographic variables are shown between the training and validation sets. **A.** Age; **B.** Sex; and **C.** Self-reported race/ethnicity. AA: African American; EA: European American: Others: All other populations (e.g., Asians).

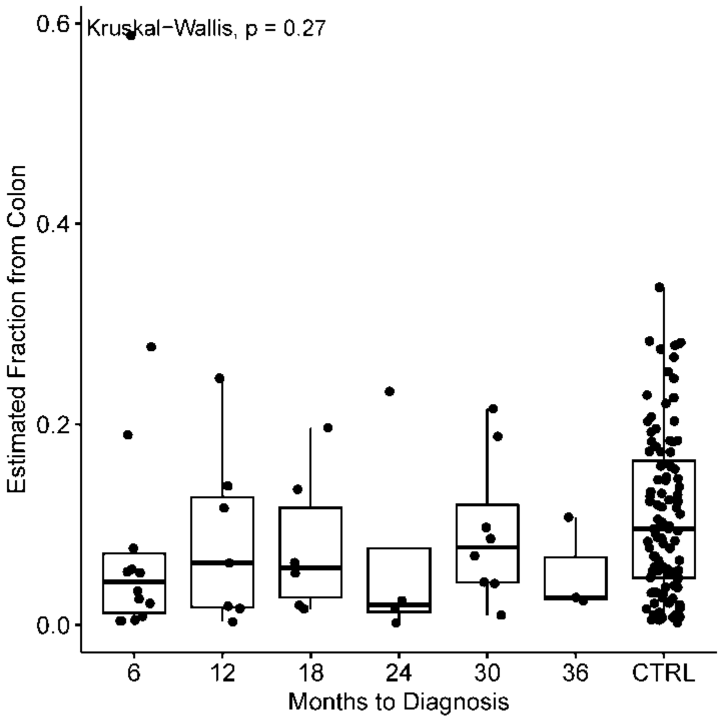


**Figure S3. Estimation of colon-derived signals through cellular deconvolution in cfDNA**. No statistically significant trend of colon-derived signals over time to diagnosis were observed.

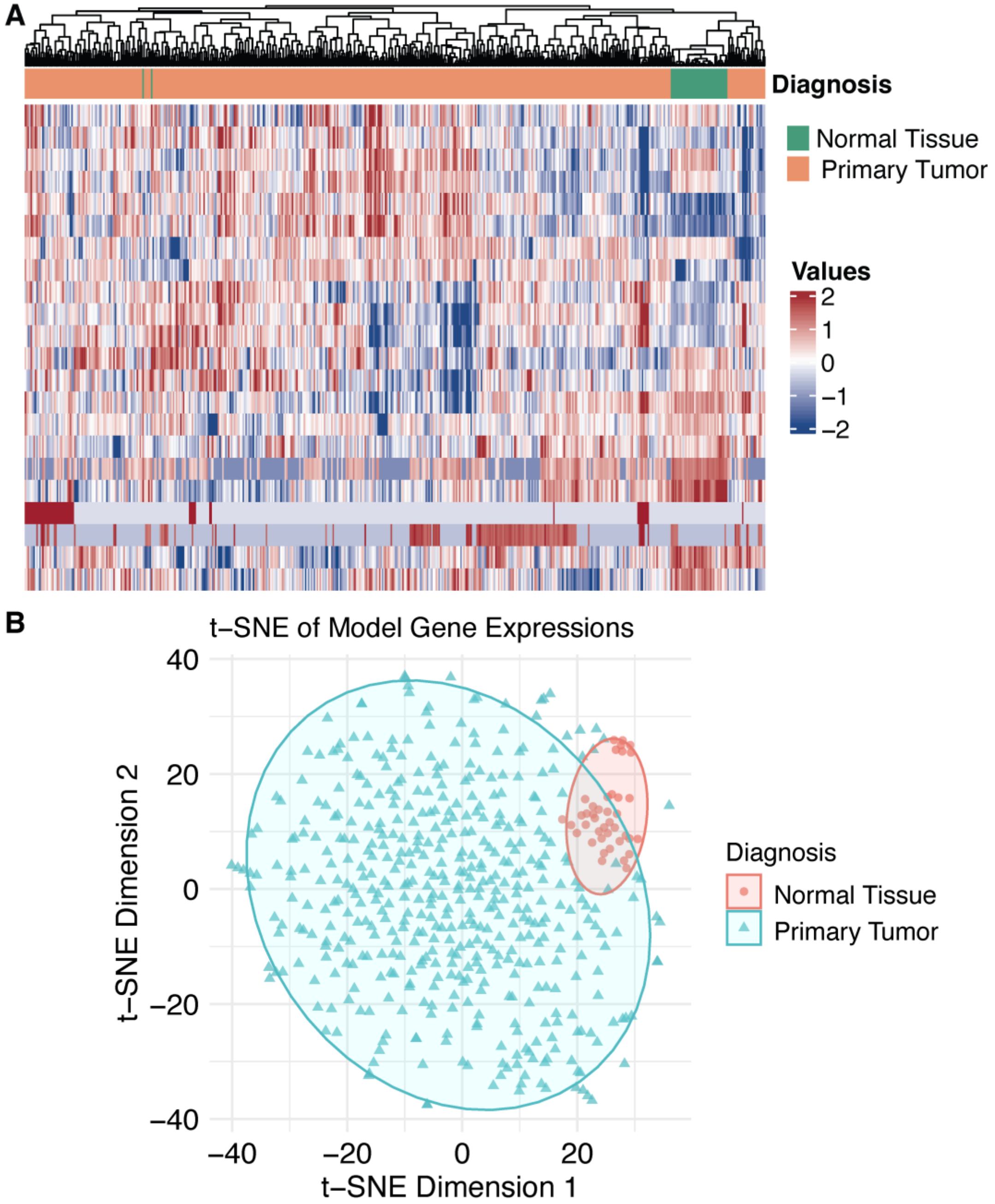


**Figure S4. Analysis of model gene expression from TCGA.**
**A.** Heatmap illustrating the expression levels of 28 model genes in TCGA dataset. **B.** t-SNE analysis depicting distinct gene expression clusters of the 28 model genes from TCGA.

**Table S1.** Multivariable Cox models identified 5hmC modification levels at 890 gene body regions are associated with pre-clinical CRC cases in the training set (p<0.05).

[Please see accompanying spreadsheet appended at the end of this document]

**Table S2.** Enriched KEGG pathways for the 540 pre-clinical CRC-associate genes.

| ID | Description | GeneRatio | BgRatio | pvalue | geneSYMBOL | Count | model_gene |
|---|---|---|---|---|---|---|---|
| hsa04810 | Regulation of actin cytoskeleton | 13/240 | 229/8622 | 0.0115522 | CHRM5, RDX, SSH3, FGF18, MYH9, RRAS2, CXCR4, ITGAE, ACTB, ARHGEF1, BRAF, RAC2, ACTN1 | 13 | |
| hsa05205 | Proteoglycans in cancer | 12/240 | 205/8622 | 0.0119629 | TGFB2, RDX, CAMK2G, FRS2, FZD5, RRAS2, ACTB, TGFB1, WNT10A, ARHGEF1, BRAF, MET | 12 | TGFB2 |
| hsa04015 | Rap1 signaling pathway | 12/240 | 210/8622 | 0.0142663 | TLN2, ADORA2B, FGF18, ACTB, RAPGEF4, ADCY4, ADCY4, PLCB2, BRAF, RAC2, CALML5, MET, KITLG | 12 | TLN2 |
| hsa04020 | Calcium signaling pathway | 13/240 | 253/8622 | 0.0243698 | CHRM5, ADORA2B, FGF18, CAMK2G, CXCR4, RYR1, P2RX6, ADCY4, ADCY4, PLCB2, PTAFR, NTRK1, CALML5, MET | 13 | |
| hsa05202 | Transcriptional misregulation in cancer | 10/240 | 193/8622 | 0.0430933 | KMT2A, PPARG, BCL2L1, RXRG, SUPT3H, PBX3, NTRK1, CDK9, MET, ZEB1 | 10 | KMT2A |

**Table S3.** List of 32 Differentially 5hmC-modified genes

| Feature gene |
|---|
| AC006372.1 |
| ACAD8 |
| BBS4 |
| BCL2L2-PABPN1 |
| BTF3L4 |
| CCL1 |
| COX5B |
| DAAM2 |
| ENSA |
| GBX1 |
| HDHD3 |
| IL10 |
| KMT2A |
| MALSU1 |
| MRPL19 |
| MYADML2 |
| NMNAT2 |
| NRL |
| PHOX2A |
| PPP1R1B |
| PRCP |
| RP11-201K10.3 |
| RP3-382I10.7 |
| RPL10A |
| SLC22A6 |
| SLC7A4 |
| TGFB2 |
| THUMPD3 |
| TIMM21 |
| TLN2 |
| TRAIP |
| ZNF624 |

**Table S4.** Model genes demonstrated relevance in gene expression based on data from TCGA.

| Gene | logFC | AveExpr | t | p-value | FDR | B |
|---|---|---|---|---|---|---|
| DAAM2 | 2.117271829 | 14.4307451 | 12.93220943 | 2.78E-33 | 7.78E-32 | 64.67868589 |
| TRAIP | -1.223337667 | 15.99913725 | -11.49690647 | 2.34E-27 | 3.27E-26 | 51.14683236 |
| MALSU1 | -0.83869988 | 17.22737255 | -9.50237549 | 8.01E-20 | 7.47E-19 | 33.97483206 |
| PHOX2A | 7.419916116 | 6.265123529 | 9.357463914 | 2.60E-19 | 1.82E-18 | 32.81284855 |
| MYADML2 | -4.09604436 | 12.87087647 | -8.087827915 | 4.47E-15 | 2.50E-14 | 23.19404953 |
| TLN2 | 1.000081127 | 15.91909804 | 7.523247519 | 2.43E-13 | 1.14E-12 | 19.26268361 |
| ACAD8 | 0.438141869 | 16.30366667 | 7.257073675 | 1.48E-12 | 5.94E-12 | 17.48799688 |
| ZNF624 | 0.852382079 | 13.49126471 | 6.599939449 | 1.03E-10 | 3.62E-10 | 13.33206752 |
| THUMPD3 | -0.323006397 | 17.34703922 | -4.930543961 | 1.11E-06 | 3.46E-06 | 4.316242192 |
| IL10 | 1.891355869 | 11.70313333 | 4.650923675 | 4.22E-06 | 1.18E-05 | 3.035586123 |

**Table S1**. Multivariable Cox models identified 5hmC modification levels at 890 gene body regions are associated with pre-clinical CRC cases in the training set ($p<0.05$).

| gene | coefficient | exp(coefficient) | standard error | z.statistics | p.value | same association direction in validation set |
|---|---|---|---|---|---|---|
| STAC2 | 1.688195697 | 5.409711138 | 0.408604149 | 4.131616627 | 3.60E-05 | TRUE |
| RAB37 | 2.33238986 | 10.30253375 | 0.620917146 | 3.756362463 | 0.000172 | TRUE |
| NRBP1 | 2.339939839 | 10.38061204 | 0.648585344 | 3.607759351 | 0.000309 | TRUE |
| NAA30 | -1.813727518 | 0.163045249 | 0.506169338 | -3.583242569 | 0.000339 | TRUE |
| MRPL19 | -2.068016508 | 0.126436319 | 0.58823214 | -3.515646918 | 0.000439 | TRUE |
| MYRIP | -2.354856246 | 0.094907149 | 0.672977085 | -3.4991626 | 0.000467 | TRUE |
| LCAT | 1.872814483 | 6.506583322 | 0.548332559 | 3.415471969 | 0.000637 | FALSE |
| CACNB1 | 1.268796837 | 3.556570852 | 0.374445348 | 3.388470025 | 0.000703 | TRUE |
| PIEZO2 | -2.264903462 | 0.103840059 | 0.678098069 | -3.340082452 | 0.000838 | FALSE |
| MYO16 | -2.217590769 | 0.108871089 | 0.664820674 | -3.335622456 | 0.000851 | FALSE |
| MEAF6 | -2.179500195 | 0.113098043 | 0.655352262 | -3.325692643 | 0.000882 | TRUE |
| NEO1 | -1.905142148 | 0.148801489 | 0.574586527 | -3.315674937 | 0.000914 | TRUE |
| MTPAP | -2.603868455 | 0.073986809 | 0.791104134 | -3.291435781 | 0.000997 | FALSE |
| STX4 | 1.674171342 | 5.334372945 | 0.51528128 | 3.249043591 | 0.001158 | TRUE |
| TLN2 | -2.248503066 | 0.105557118 | 0.696833083 | -3.226745573 | 0.001252 | TRUE |
| AARD | -1.61979296 | 0.197939676 | 0.505999019 | -3.201178067 | 0.001369 | FALSE |
| PGAP3 | 0.89317534 | 2.442874305 | 0.280905229 | 3.179632299 | 0.001475 | TRUE |
| TBC1D20 | 2.601027584 | 13.47758026 | 0.82559765 | 3.150478425 | 0.00163 | TRUE |
| PLPPR2 | 1.21922399 | 3.384560262 | 0.38780205 | 3.143933842 | 0.001667 | TRUE |
| TSPAN12 | -1.630089927 | 0.195911956 | 0.519740126 | -3.136355737 | 0.001711 | TRUE |
| TMEM26 | -1.808820097 | 0.163847347 | 0.580987311 | -3.113355599 | 0.00185 | FALSE |
| ENPP6 | -2.376611208 | 0.092864744 | 0.768065026 | -3.094283853 | 0.001973 | TRUE |
| RTCA | -2.000970122 | 0.135204055 | 0.648480321 | -3.085629673 | 0.002031 | TRUE |
| WIPF2 | 1.165100376 | 3.206244697 | 0.379130621 | 3.073084342 | 0.002119 | FALSE |
| PADI1 | 1.869688046 | 6.486272666 | 0.610749276 | 3.061302106 | 0.002204 | TRUE |
| KMT2A | -2.386989145 | 0.091905983 | 0.78062191 | -3.057804441 | 0.00223 | TRUE |
| PSD3 | -1.686334612 | 0.185197101 | 0.55149784 | -3.057735658 | 0.00223 | FALSE |
| RNPC3 | -1.701968466 | 0.182324271 | 0.557206795 | -3.054464663 | 0.002255 | FALSE |
| CCDC122 | -1.872241933 | 0.153778514 | 0.613269171 | -3.052887739 | 0.002267 | FALSE |
| C17orf99 | 1.837311257 | 6.279631225 | 0.603160357 | 3.046140609 | 0.002318 | TRUE |
| LONRF1 | -1.817353922 | 0.162455051 | 0.596719784 | -3.045573432 | 0.002322 | TRUE |
| NUDCD2 | -1.767815098 | 0.170705557 | 0.581358673 | -3.040833792 | 0.002359 | FALSE |
| SLC46A1 | 1.710834259 | 5.533575985 | 0.562789535 | 3.039918392 | 0.002366 | TRUE |
| CD40 | 1.989601006 | 7.312615487 | 0.65893364 | 3.019425455 | 0.002533 | FALSE |
| SPESP1 | -2.207111025 | 0.110018029 | 0.732595446 | -3.012728289 | 0.002589 | FALSE |
| GUCD1 | 1.792247335 | 6.002927911 | 0.598006957 | 2.997034257 | 0.002726 | TRUE |
| BRF2 | -1.59296717 | 0.203321427 | 0.531993305 | -2.994336877 | 0.00275 | TRUE |

| | | | | | | |
|---|---|---|---|---|---|---|
| TIMM21 | -1.977853558 | 0.138365913 | 0.664839178 | -2.974935326 | 0.002931 | TRUE |
| DDIT4 | 1.380163844 | 3.975552943 | 0.464557563 | 2.970921054 | 0.002969 | FALSE |
| PMS2 | -1.865309665 | 0.154848251 | 0.630604496 | -2.957970768 | 0.003097 | TRUE |
| LYPLA2 | 1.345802175 | 3.841266673 | 0.455075911 | 2.95731359 | 0.003103 | TRUE |
| CTC-554D6.1 | -1.816887461 | 0.162530848 | 0.614841368 | -2.955050775 | 0.003126 | FALSE |
| C1QTNF1 | 1.491643471 | 4.444393753 | 0.506978842 | 2.942220362 | 0.003259 | TRUE |
| LRRC2 | -2.051413955 | 0.128553007 | 0.69988183 | -2.931086175 | 0.003378 | FALSE |
| SOGA1 | 2.014643768 | 7.498055861 | 0.687804429 | 2.929093915 | 0.0034 | TRUE |
| ACAD8 | 1.949266335 | 7.023532767 | 0.665995168 | 2.926847564 | 0.003424 | TRUE |
| C12orf76 | 1.99904489 | 7.382002108 | 0.683316076 | 2.92550543 | 0.003439 | FALSE |
| RP11-201K10.3 | 1.360847909 | 3.899498319 | 0.465722658 | 2.922013535 | 0.003478 | TRUE |
| GALT | 1.68377402 | 5.385843944 | 0.577022087 | 2.918040849 | 0.003522 | TRUE |
| C9orf84 | -1.712030408 | 0.180498934 | 0.587098013 | -2.916089597 | 0.003544 | FALSE |
| NIT1 | 1.608481681 | 4.995221129 | 0.551798533 | 2.914979987 | 0.003557 | TRUE |
| CATSPERG | 1.693849038 | 5.440380691 | 0.582405843 | 2.908365462 | 0.003633 | FALSE |
| EIF2B4 | 1.731103442 | 5.646881478 | 0.596477892 | 2.90220889 | 0.003705 | FALSE |
| RP3-382I10.7 | -1.598695696 | 0.202160024 | 0.550903267 | -2.901953561 | 0.003708 | TRUE |
| R3HCC1 | -1.852003887 | 0.156922396 | 0.644874467 | -2.871882796 | 0.00408 | FALSE |
| RP11-101E3.5 | -1.601890475 | 0.201515198 | 0.55790998 | -2.871234668 | 0.004089 | FALSE |
| AQR | -1.968694475 | 0.139639039 | 0.690004959 | -2.853159895 | 0.004329 | FALSE |
| C1orf131 | -1.822857689 | 0.161563393 | 0.638940886 | -2.852936366 | 0.004332 | TRUE |
| PTPRG | -1.401841607 | 0.246143247 | 0.492012306 | -2.849200295 | 0.004383 | TRUE |
| BACH2 | -2.171716754 | 0.11398177 | 0.763109791 | -2.845877198 | 0.004429 | TRUE |
| UBA2 | -1.781228682 | 0.168431072 | 0.625902625 | -2.845855905 | 0.004429 | TRUE |
| HERC4 | -1.839747256 | 0.158857571 | 0.648346919 | -2.837596974 | 0.004545 | FALSE |
| SH3BGR | -1.916786638 | 0.147078821 | 0.676213141 | -2.834589454 | 0.004588 | TRUE |
| RNF14 | -1.631425678 | 0.195650441 | 0.57626391 | -2.831039129 | 0.00464 | FALSE |
| MADD | 2.430267322 | 11.36191897 | 0.861982291 | 2.819393562 | 0.004811 | TRUE |
| FANCL | -1.433786032 | 0.238404604 | 0.508656938 | -2.81876826 | 0.004821 | TRUE |
| SPNS1 | 1.482982716 | 4.406068149 | 0.527497871 | 2.811352987 | 0.004933 | FALSE |
| GTF2E2 | -1.890291798 | 0.151027733 | 0.672992172 | -2.808787199 | 0.004973 | TRUE |
| RNF152 | -1.446806015 | 0.2353207 | 0.515286744 | -2.807768744 | 0.004989 | TRUE |
| AMBRA1 | -3.220562434 | 0.039932592 | 1.148860196 | -2.803267487 | 0.005059 | TRUE |
| C4BPB | -1.563980032 | 0.209301385 | 0.558036629 | -2.802647624 | 0.005069 | TRUE |
| HSD11B2 | 1.447426748 | 4.252158551 | 0.517904264 | 2.794776656 | 0.005194 | FALSE |
| C11orf71 | -1.582410244 | 0.205479246 | 0.566401064 | -2.793798151 | 0.005209 | FALSE |
| CHRM5 | 1.749232053 | 5.75018514 | 0.626493207 | 2.792100588 | 0.005237 | TRUE |
| LYRM5 | -1.454613007 | 0.233490706 | 0.521955939 | -2.786850191 | 0.005322 | FALSE |
| CCS | 1.399333509 | 4.052498113 | 0.502744972 | 2.783386381 | 0.005379 | FALSE |
| AQP2 | 1.538913857 | 4.659526607 | 0.553077008 | 2.78245856 | 0.005395 | FALSE |
| PAXIP1 | -2.167916443 | 0.11441576 | 0.781867576 | -2.772741204 | 0.005559 | TRUE |
| SPEM1 | 1.350846124 | 3.860690773 | 0.487203598 | 2.772652191 | 0.00556 | FALSE |
| HSP90B1 | -1.795329462 | 0.166072729 | 0.648465504 | -2.768581293 | 0.00563 | TRUE |
| SELM | 1.383821166 | 3.990119443 | 0.500593868 | 2.764359005 | 0.005703 | FALSE |

| | | | | | | |
|---|---|---|---|---|---|---|
| EPS8L1 | 1.26839448 | 3.555140131 | 0.459492631 | 2.760423984 | 0.005773 | FALSE |
| NRBF2 | 1.746158596 | 5.732539327 | 0.634916924 | 2.750215865 | 0.005956 | FALSE |
| TMC4 | 1.358497351 | 3.890343089 | 0.494137906 | 2.749227156 | 0.005974 | TRUE |
| CLEC2A | -1.515818499 | 0.219628346 | 0.551421108 | -2.748930856 | 0.005979 | FALSE |
| SYCE3 | 1.717871439 | 5.572654094 | 0.625025038 | 2.748484195 | 0.005987 | FALSE |
| NRN1 | 1.385175053 | 3.995525272 | 0.504379356 | 2.746296088 | 0.006027 | TRUE |
| PENK | 1.533300223 | 4.633443012 | 0.559273104 | 2.741594779 | 0.006114 | FALSE |
| DEDD2 | 1.76920292 | 5.866175687 | 0.645360024 | 2.741420067 | 0.006117 | FALSE |
| ROPN1L | 1.593757022 | 4.922207072 | 0.581403944 | 2.741221554 | 0.006121 | TRUE |
| TMEM128 | -1.706990511 | 0.181410926 | 0.623671153 | -2.737004115 | 0.0062 | TRUE |
| C9orf66 | 1.508211693 | 4.518642846 | 0.551384697 | 2.73531656 | 0.006232 | FALSE |
| PPP1R27 | 1.462633066 | 4.317312344 | 0.535017629 | 2.7338035 | 0.006261 | FALSE |
| TUBGCP5 | -1.663239607 | 0.189524001 | 0.60908162 | -2.730733537 | 0.006319 | TRUE |
| C10orf35 | -1.440797875 | 0.236738795 | 0.528111433 | -2.728208074 | 0.006368 | TRUE |
| NR1D2 | -1.713568334 | 0.180221553 | 0.628937821 | -2.724543311 | 0.006439 | FALSE |
| ADGRE5 | 1.385474338 | 3.996721253 | 0.508530737 | 2.72446528 | 0.006441 | TRUE |
| QRFP | 1.445132811 | 4.242415545 | 0.530930267 | 2.721888164 | 0.006491 | TRUE |
| RECQL4 | 0.959971126 | 2.611621064 | 0.35303043 | 2.719230537 | 0.006543 | TRUE |
| AGAP1 | -1.145824784 | 0.31796156 | 0.421460238 | -2.718701979 | 0.006554 | TRUE |
| PDCD4 | -1.359267677 | 0.256848804 | 0.500279516 | -2.717016453 | 0.006587 | FALSE |
| CCL1 | 1.63659529 | 5.1376475 | 0.603459118 | 2.712023469 | 0.006687 | TRUE |
| RACGAP1 | 2.130010857 | 8.414958168 | 0.787435343 | 2.704997782 | 0.00683 | TRUE |
| CEP290 | -1.379282358 | 0.251759161 | 0.510500593 | -2.701823225 | 0.006896 | TRUE |
| SEC31A | -1.866936923 | 0.154596478 | 0.69100921 | -2.701754037 | 0.006897 | TRUE |
| ARHGAP27 | 1.140005425 | 3.126785328 | 0.422338083 | 2.699272147 | 0.006949 | FALSE |
| SSC5D | 1.355872801 | 3.880146074 | 0.502960561 | 2.695783539 | 0.007022 | FALSE |
| NADK2 | -1.513549896 | 0.220127161 | 0.561690663 | -2.694632465 | 0.007047 | TRUE |
| ARL11 | 1.655415216 | 5.235253231 | 0.617834126 | 2.679384558 | 0.007376 | FALSE |
| HELQ | -1.861777927 | 0.155396102 | 0.695103796 | -2.678417148 | 0.007397 | TRUE |
| PAQR7 | 1.441195491 | 4.225744637 | 0.538289189 | 2.677362876 | 0.00742 | FALSE |
| ADORA2B | 1.727792633 | 5.628216646 | 0.645369411 | 2.677214947 | 0.007424 | TRUE |
| PRCP | -1.691565144 | 0.18423095 | 0.633190106 | -2.671496488 | 0.007551 | TRUE |
| TGFB2 | -1.554422992 | 0.211311276 | 0.581905861 | -2.671261962 | 0.007557 | TRUE |
| CD59 | -1.782104491 | 0.168283623 | 0.667286863 | -2.670672224 | 0.00757 | TRUE |
| VEGFB | 1.419088465 | 4.133351028 | 0.531479377 | 2.670072495 | 0.007583 | FALSE |
| COMMD10 | -1.601116344 | 0.201671258 | 0.59968832 | -2.669914173 | 0.007587 | TRUE |
| MRPL3 | -1.503356405 | 0.2223825 | 0.563178917 | -2.669411727 | 0.007598 | FALSE |
| AP1M2 | 1.466248191 | 4.332948213 | 0.550069481 | 2.665569065 | 0.007686 | TRUE |
| CCDC155 | 1.412608419 | 4.106653319 | 0.530052756 | 2.665033625 | 0.007698 | TRUE |
| PATL1 | -1.608893717 | 0.200108869 | 0.604019292 | -2.663646243 | 0.00773 | TRUE |
| TPRA1 | 1.598537759 | 4.945795189 | 0.602593471 | 2.652763155 | 0.007984 | TRUE |
| RDX | -2.068762264 | 0.126342063 | 0.780882085 | -2.649263317 | 0.008067 | TRUE |
| C11orf74 | -1.565963714 | 0.208886609 | 0.591307909 | -2.648305038 | 0.00809 | FALSE |
| ARHGEF26 | -1.759081865 | 0.172202897 | 0.664296146 | -2.648038642 | 0.008096 | FALSE |

| | | | | | | |
|---|---|---|---|---|---|---|
| MED13 | -1.672767937 | 0.18772673 | 0.632384185 | -2.645176742 | 0.008165 | TRUE |
| ALCAM | -1.320934412 | 0.266885804 | 0.50024771 | -2.640560636 | 0.008277 | FALSE |
| DNAJC8 | -1.675266836 | 0.187258206 | 0.634459396 | -2.640463435 | 0.008279 | FALSE |
| AKAP11 | -1.513233869 | 0.220196738 | 0.573170461 | -2.640111403 | 0.008288 | TRUE |
| LCLAT1 | -1.573631995 | 0.207290934 | 0.596052047 | -2.640091588 | 0.008288 | TRUE |
| ZNF460 | -1.273210082 | 0.279931575 | 0.482369158 | -2.639493136 | 0.008303 | FALSE |
| PPP4R2 | -1.780009301 | 0.168636579 | 0.674609431 | -2.638577554 | 0.008325 | FALSE |
| MYH10 | -1.951211036 | 0.142101877 | 0.741987009 | -2.629710509 | 0.008546 | FALSE |
| TMEM150B | 1.650014034 | 5.207052902 | 0.62804888 | 2.627206393 | 0.008609 | TRUE |
| FAM134B | -1.235667885 | 0.290640583 | 0.470376364 | -2.626976992 | 0.008615 | TRUE |
| CFAP58 | -1.562440043 | 0.209623955 | 0.596215511 | -2.6205961 | 0.008778 | FALSE |
| SRP54 | -1.683592961 | 0.185705543 | 0.643467184 | -2.616439506 | 0.008885 | TRUE |
| GNPNAT1 | -1.559508131 | 0.210239456 | 0.596055621 | -2.616380209 | 0.008887 | TRUE |
| TFB1M | -1.534265241 | 0.215614057 | 0.587200478 | -2.612847399 | 0.008979 | TRUE |
| KCNF1 | 1.170355095 | 3.223136956 | 0.448660036 | 2.608556595 | 0.009092 | TRUE |
| NFE2 | 1.339231271 | 3.816108824 | 0.513410436 | 2.60850029 | 0.009094 | FALSE |
| NAA35 | -1.59009953 | 0.203905316 | 0.609597388 | -2.608442166 | 0.009096 | TRUE |
| LACTB2 | -1.603836721 | 0.201123382 | 0.61532242 | -2.606498102 | 0.009147 | FALSE |
| HS2ST1 | -1.622585318 | 0.197387729 | 0.622787405 | -2.60535988 | 0.009178 | TRUE |
| RP11-12J10.3 | 1.480181684 | 4.39374388 | 0.570243134 | 2.595702772 | 0.00944 | FALSE |
| FHL3 | 1.437118971 | 4.208553368 | 0.553960822 | 2.594261024 | 0.009479 | TRUE |
| EED | -1.572263824 | 0.207574737 | 0.60854707 | -2.583635519 | 0.009777 | TRUE |
| TFDP2 | -2.34034693 | 0.096294225 | 0.906420228 | -2.581966793 | 0.009824 | FALSE |
| SMAD4 | -1.481459584 | 0.227305675 | 0.573774271 | -2.581955409 | 0.009824 | FALSE |
| UPF2 | -1.954644455 | 0.141614818 | 0.757306715 | -2.581047304 | 0.00985 | TRUE |
| SVEP1 | -1.493316624 | 0.224626417 | 0.578811259 | -2.579971626 | 0.009881 | TRUE |
| FABP6 | 1.304362099 | 3.685337464 | 0.505773703 | 2.578944085 | 0.00991 | TRUE |
| MED10 | -1.587465374 | 0.204443143 | 0.615777972 | -2.577983375 | 0.009938 | TRUE |
| BBS4 | -1.84896359 | 0.157400213 | 0.717857312 | -2.575670068 | 0.010005 | TRUE |
| PHF19 | 1.945501824 | 6.997142305 | 0.758134824 | 2.566168656 | 0.010283 | TRUE |
| TMA16 | -1.38942408 | 0.249218793 | 0.541967466 | -2.563666949 | 0.010357 | TRUE |
| KCNMA1 | -2.002878025 | 0.134946345 | 0.781492487 | -2.562888393 | 0.010381 | TRUE |
| SLC25A25 | 1.75255977 | 5.769352003 | 0.68383847 | 2.562827109 | 0.010382 | FALSE |
| GZF1 | 1.62225261 | 5.064485789 | 0.633015216 | 2.562738729 | 0.010385 | FALSE |
| SFTPB | 1.389224661 | 4.011738388 | 0.54213429 | 2.562510221 | 0.010392 | TRUE |
| BLZF1 | -1.436976272 | 0.237645248 | 0.561024339 | -2.561343907 | 0.010427 | TRUE |
| AP1G2 | 1.534028998 | 4.636820983 | 0.599304573 | 2.559681784 | 0.010477 | FALSE |
| TAGLN2 | 1.446884071 | 4.249851626 | 0.56611199 | 2.55582658 | 0.010594 | TRUE |
| TNRC6A | -1.788136131 | 0.167271652 | 0.699901523 | -2.554839604 | 0.010624 | TRUE |
| SSH3 | 1.088273891 | 2.969144579 | 0.426037413 | 2.554409209 | 0.010637 | TRUE |
| ANGPTL6 | 1.515431478 | 4.551384526 | 0.593482349 | 2.553456695 | 0.010666 | TRUE |
| RPSA | 1.494960486 | 4.459160348 | 0.585790787 | 2.552038234 | 0.010709 | FALSE |
| ZCCHC8 | -1.819336815 | 0.16213324 | 0.712931388 | -2.551910108 | 0.010713 | FALSE |
| RARA | 0.912107614 | 2.489564048 | 0.357422136 | 2.551905781 | 0.010714 | FALSE |

| | | | | | | |
|---|---|---|---|---|---|---|
| HSD17B1 | 1.149205256 | 3.155683951 | 0.450435332 | 2.551321296 | 0.010732 | TRUE |
| CMC1 | -1.499945006 | 0.223142431 | 0.58847426 | -2.548871053 | 0.010807 | FALSE |
| PRMT1 | 1.252002307 | 3.497338696 | 0.491343339 | 2.548121055 | 0.01083 | FALSE |
| ZNF624 | -1.577001676 | 0.206593605 | 0.619322673 | -2.546332866 | 0.010886 | TRUE |
| FCAMR | 1.762137842 | 5.824876757 | 0.692524964 | 2.544511656 | 0.010943 | TRUE |
| SDAD1 | -1.725745565 | 0.178040262 | 0.67824733 | -2.544419253 | 0.010946 | TRUE |
| TOR1AIP2 | -1.505978964 | 0.221800053 | 0.592138255 | -2.543289428 | 0.010981 | TRUE |
| METAP1D | -1.591978002 | 0.203522645 | 0.626216183 | -2.542217921 | 0.011015 | TRUE |
| UBXN2B | -1.549691787 | 0.212313402 | 0.609619737 | -2.542063017 | 0.01102 | FALSE |
| FZD6 | -1.498256945 | 0.223519427 | 0.590366442 | -2.537842325 | 0.011154 | FALSE |
| FGF18 | 1.195195652 | 3.304204182 | 0.471131292 | 2.536863231 | 0.011185 | TRUE |
| DAPL1 | -1.588514399 | 0.204228789 | 0.626364496 | -2.536086272 | 0.01121 | TRUE |
| SLC14A1 | -1.578710315 | 0.206240913 | 0.622504912 | -2.536060816 | 0.011211 | FALSE |
| TGFB1I1 | 1.094520711 | 2.987750344 | 0.432190748 | 2.532494544 | 0.011325 | FALSE |
| SERTAD4 | -1.560127457 | 0.21010929 | 0.616108153 | -2.532229852 | 0.011334 | TRUE |
| RNF214 | -1.996536757 | 0.135804795 | 0.788950828 | -2.53062255 | 0.011386 | TRUE |
| KCTD19 | 1.861239436 | 6.431703515 | 0.736151774 | 2.528336547 | 0.01146 | FALSE |
| FUCA1 | -1.646135972 | 0.19279343 | 0.651304543 | -2.527444328 | 0.01149 | FALSE |
| SYNGAP1 | 1.640853855 | 5.159573159 | 0.649458063 | 2.526497013 | 0.011521 | TRUE |
| BTBD9 | -2.427538478 | 0.088253804 | 0.961749946 | -2.524084861 | 0.0116 | FALSE |
| URAD | -1.614422275 | 0.199005608 | 0.640554209 | -2.5203523 | 0.011724 | FALSE |
| SWI5 | 1.23067839 | 3.423551251 | 0.488433048 | 2.519646027 | 0.011747 | TRUE |
| SWSAP1 | 1.41216529 | 4.104833946 | 0.560580289 | 2.519113352 | 0.011765 | TRUE |
| FCHO1 | 1.483334428 | 4.407618089 | 0.589257713 | 2.517293191 | 0.011826 | TRUE |
| MICU3 | -1.27978533 | 0.278096993 | 0.508603919 | -2.516271076 | 0.01186 | TRUE |
| DECR2 | 1.102968516 | 3.013097187 | 0.438643811 | 2.514496928 | 0.01192 | TRUE |
| ZFP90 | -1.676733611 | 0.186983741 | 0.667247755 | -2.512910082 | 0.011974 | TRUE |
| IRF3 | 1.306520608 | 3.693300889 | 0.520006772 | 2.512506912 | 0.011988 | FALSE |
| PGGT1B | -1.283624223 | 0.277031455 | 0.510903621 | -2.512458653 | 0.011989 | TRUE |
| PLEKHB2 | 2.224982178 | 9.253317887 | 0.887081102 | 2.508206039 | 0.012135 | TRUE |
| FAM150B | -1.439538245 | 0.237037186 | 0.574165594 | -2.50718305 | 0.01217 | TRUE |
| TOM1L2 | 2.325429132 | 10.23106962 | 0.927622286 | 2.506870702 | 0.012181 | TRUE |
| TMX4 | -1.552323692 | 0.211755348 | 0.619355169 | -2.506354623 | 0.012198 | TRUE |
| KIF21B | 1.187516069 | 3.278926457 | 0.474236191 | 2.504060408 | 0.012278 | FALSE |
| RAP1GAP2 | 1.362957144 | 3.907731957 | 0.544315381 | 2.503984252 | 0.01228 | TRUE |
| AFF4 | -1.791036319 | 0.166787235 | 0.716823087 | -2.498575104 | 0.012469 | TRUE |
| COQ7 | -1.369781376 | 0.25416252 | 0.548252294 | -2.498450789 | 0.012474 | TRUE |
| SLC7A4 | 1.16219664 | 3.196948111 | 0.465648708 | 2.495865701 | 0.012565 | TRUE |
| THRA | 1.025674878 | 2.788977045 | 0.411363528 | 2.493353951 | 0.012654 | FALSE |
| C11orf68 | 1.193901746 | 3.299931617 | 0.478886734 | 2.493077512 | 0.012664 | TRUE |
| DICER1 | -1.398544376 | 0.246956178 | 0.561096855 | -2.492518649 | 0.012684 | TRUE |
| POU2F2 | 1.613995255 | 5.022838714 | 0.647641965 | 2.492110368 | 0.012699 | FALSE |
| FBXO39 | 1.537596487 | 4.653392331 | 0.618235123 | 2.48707398 | 0.01288 | FALSE |
| NOD1 | 1.980145535 | 7.243797137 | 0.797536564 | 2.482827276 | 0.013034 | FALSE |

| | | | | | | |
|---|---|---|---|---|---|---|
| MOCS3 | 1.488786534 | 4.431714521 | 0.599705783 | 2.482528229 | 0.013045 | FALSE |
| ZNF682 | -1.30839039 | 0.270254711 | 0.527723518 | -2.479310368 | 0.013164 | TRUE |
| CCDC140 | 1.269892918 | 3.560471278 | 0.512711792 | 2.476816287 | 0.013256 | FALSE |
| CPAMD8 | 1.48890067 | 4.432220367 | 0.601236669 | 2.476396979 | 0.013272 | TRUE |
| TMEM57 | -1.793320417 | 0.166406712 | 0.725374253 | -2.472269188 | 0.013426 | TRUE |
| ZFYVE26 | -1.889328888 | 0.151173229 | 0.764361897 | -2.471772724 | 0.013444 | FALSE |
| DNTTIP2 | -1.487705748 | 0.225890311 | 0.602256994 | -2.470217472 | 0.013503 | FALSE |
| FCER2 | 1.44648115 | 4.248139618 | 0.585644623 | 2.469895722 | 0.013515 | FALSE |
| SLC39A9 | -1.524703836 | 0.217685518 | 0.618242182 | -2.466191858 | 0.013656 | TRUE |
| SLC16A1 | -1.585882808 | 0.204766943 | 0.643265303 | -2.465363514 | 0.013687 | TRUE |
| RITA1 | 1.407970759 | 4.087652153 | 0.571150023 | 2.46515049 | 0.013696 | TRUE |
| PNPT1 | -1.676569789 | 0.187014376 | 0.680222829 | -2.46473614 | 0.013711 | FALSE |
| USP4 | 1.844471428 | 6.324755817 | 0.748420728 | 2.464484692 | 0.013721 | TRUE |
| ZNF543 | 1.476275462 | 4.37661442 | 0.599655823 | 2.461871303 | 0.013821 | TRUE |
| SEPT2 | -1.582009243 | 0.20556166 | 0.643037947 | -2.460211331 | 0.013886 | FALSE |
| MRPL57 | -1.309499263 | 0.269955199 | 0.53323409 | -2.455768089 | 0.014058 | TRUE |
| MRPS9 | -1.395697928 | 0.247660127 | 0.568474034 | -2.455165661 | 0.014082 | FALSE |
| MPV17 | 1.549822465 | 4.710633806 | 0.631356374 | 2.454750643 | 0.014098 | TRUE |
| MYCL | 1.515653799 | 4.552396507 | 0.617616777 | 2.45403599 | 0.014126 | FALSE |
| GIN1 | -1.418337836 | 0.242116119 | 0.57842418 | -2.452072175 | 0.014204 | FALSE |
| ADAM23 | -1.234699654 | 0.290922127 | 0.503654985 | -2.45147907 | 0.014227 | FALSE |
| SULT6B1 | 1.4773889 | 4.381490224 | 0.603647804 | 2.447435227 | 0.014388 | FALSE |
| MYO1A | 1.62562388 | 5.081588352 | 0.664228559 | 2.447386306 | 0.01439 | FALSE |
| DAPK2 | 1.913791278 | 6.778740228 | 0.782014414 | 2.447258315 | 0.014395 | TRUE |
| PRSS12 | -1.507403803 | 0.221484249 | 0.61641692 | -2.445428984 | 0.014468 | TRUE |
| GNAI1 | -1.098987862 | 0.333208166 | 0.449605352 | -2.444338924 | 0.014512 | FALSE |
| LFNG | 0.758268408 | 2.134576802 | 0.310452113 | 2.442464962 | 0.014587 | TRUE |
| TACR2 | 1.438864951 | 4.215907838 | 0.589159779 | 2.442232143 | 0.014597 | FALSE |
| APOBEC3A | 0.900404003 | 2.460596999 | 0.368745757 | 2.441801667 | 0.014614 | TRUE |
| NIPBL | -1.263574358 | 0.282641955 | 0.517584326 | -2.441291775 | 0.014635 | TRUE |
| RFC1 | -1.553929684 | 0.211415543 | 0.637209855 | -2.438646659 | 0.014742 | TRUE |
| CD300C | 1.192440695 | 3.295113767 | 0.489416798 | 2.436452322 | 0.014832 | TRUE |
| RP11-540D14.8 | 1.384216458 | 3.991697018 | 0.568252529 | 2.435917814 | 0.014854 | TRUE |
| PDSS2 | -1.774219081 | 0.169615854 | 0.728718312 | -2.434711811 | 0.014904 | TRUE |
| C4orf26 | 1.611465624 | 5.010148844 | 0.662133825 | 2.433746113 | 0.014943 | FALSE |
| FBXW2 | -1.36908278 | 0.254340138 | 0.562620501 | -2.433403649 | 0.014958 | TRUE |
| GTPBP10 | -1.430518371 | 0.239184903 | 0.5879709 | -2.432974782 | 0.014975 | FALSE |
| TPTE | 1.060983764 | 2.889211894 | 0.43610995 | 2.43283549 | 0.014981 | TRUE |
| AKR1B1 | 1.456609709 | 4.291385791 | 0.598973379 | 2.431843818 | 0.015022 | FALSE |
| TPSD1 | 1.187816551 | 3.279911864 | 0.488741813 | 2.430355903 | 0.015084 | TRUE |
| MYCBPAP | 1.405802079 | 4.078796948 | 0.578670761 | 2.429364283 | 0.015125 | TRUE |
| BCKDHB | -1.329358072 | 0.264647091 | 0.547236615 | -2.429219896 | 0.015131 | TRUE |
| STK10 | 1.597860466 | 4.94244657 | 0.657821289 | 2.429019087 | 0.01514 | TRUE |
| PLEKHG1 | -1.531230693 | 0.216269342 | 0.63081206 | -2.427396035 | 0.015208 | FALSE |

| | | | | | | |
|---|---|---|---|---|---|---|
| LRIT3 | -1.444851313 | 0.235781132 | 0.595660657 | -2.425628241 | 0.015282 | FALSE |
| KLHL7 | -1.543517385 | 0.213628365 | 0.636869241 | -2.423601716 | 0.015367 | FALSE |
| SLC25A26 | -1.766796611 | 0.170879507 | 0.729091147 | -2.42328633 | 0.015381 | TRUE |
| PNMT | 1.008160898 | 2.740556216 | 0.416349437 | 2.421429712 | 0.01546 | FALSE |
| MYH9 | 1.369887963 | 3.934909815 | 0.566807443 | 2.416848929 | 0.015656 | TRUE |
| PCNX | -1.36131127 | 0.256324446 | 0.563274474 | -2.416781395 | 0.015658 | TRUE |
| C1QTNF9B | 1.363615981 | 3.910307366 | 0.564718997 | 2.414680555 | 0.015749 | TRUE |
| PADI3 | 1.417478041 | 4.126699936 | 0.587118235 | 2.41429742 | 0.015766 | TRUE |
| LYSMD3 | 1.276573485 | 3.584336877 | 0.52932911 | 2.411681999 | 0.015879 | TRUE |
| BUB1 | -1.287907774 | 0.275847315 | 0.534269182 | -2.410597161 | 0.015926 | TRUE |
| PPM1F | 1.101371664 | 3.008289557 | 0.45693784 | 2.410331489 | 0.015938 | TRUE |
| TBCA | -1.742931346 | 0.175006643 | 0.723174403 | -2.410112055 | 0.015948 | TRUE |
| NFXL1 | -1.546425787 | 0.213007951 | 0.641677938 | -2.409971883 | 0.015954 | TRUE |
| GUK1 | 0.98990873 | 2.690988854 | 0.410847919 | 2.409428609 | 0.015978 | TRUE |
| EFCAB3 | 1.306335213 | 3.692616233 | 0.542424377 | 2.408326889 | 0.016026 | TRUE |
| IL17RB | -1.463067322 | 0.231525023 | 0.608011382 | -2.40631568 | 0.016114 | TRUE |
| SUPT4H1 | 1.545567989 | 4.690635098 | 0.642491351 | 2.405585674 | 0.016147 | TRUE |
| ALMS1 | -1.473005743 | 0.229235426 | 0.612704659 | -2.404104037 | 0.016212 | TRUE |
| MPPED2 | -1.230047589 | 0.292278668 | 0.511860573 | -2.403091105 | 0.016257 | TRUE |
| TIMM22 | 1.549718261 | 4.710142966 | 0.646383106 | 2.397522843 | 0.016506 | TRUE |
| FGB | -1.279412051 | 0.27820082 | 0.534035061 | -2.395745418 | 0.016587 | TRUE |
| GIP | 1.461096358 | 4.310682989 | 0.610112374 | 2.39479876 | 0.016629 | FALSE |
| RALGPS2 | -1.269590964 | 0.280946515 | 0.530409712 | -2.393604294 | 0.016684 | TRUE |
| RCOR3 | -1.452336827 | 0.234022778 | 0.607272178 | -2.391574781 | 0.016776 | TRUE |
| SNAPC1 | -1.623307587 | 0.197245213 | 0.679046253 | -2.39056998 | 0.016822 | TRUE |
| VEZT | -1.467376524 | 0.230529481 | 0.613940702 | -2.390094871 | 0.016844 | TRUE |
| PRR35 | 0.774683972 | 2.169906268 | 0.324392603 | 2.388106154 | 0.016935 | FALSE |
| IMPDH2 | 1.468377862 | 4.342185802 | 0.614986554 | 2.387658483 | 0.016956 | FALSE |
| TOR1B | 1.39824582 | 4.048092651 | 0.585615956 | 2.387649799 | 0.016956 | TRUE |
| ERRFI1 | 1.743498304 | 5.717309368 | 0.730406803 | 2.387023638 | 0.016985 | FALSE |
| AKAP9 | -1.367558879 | 0.254728023 | 0.572929827 | -2.386957033 | 0.016988 | FALSE |
| ERCC6L2 | -1.43672597 | 0.237704739 | 0.601937726 | -2.386834895 | 0.016994 | TRUE |
| RAB44 | 1.590945289 | 4.908386579 | 0.667549874 | 2.383260565 | 0.01716 | TRUE |
| CEP295 | -1.361958268 | 0.256158658 | 0.572356278 | -2.379563777 | 0.017333 | TRUE |
| CHRNA10 | 1.174581988 | 3.236789643 | 0.493624659 | 2.379504278 | 0.017336 | FALSE |
| BMS1 | -1.287974627 | 0.275828874 | 0.541758942 | -2.377394311 | 0.017435 | TRUE |
| GLYATL3 | -1.233691137 | 0.291215674 | 0.518986461 | -2.377116225 | 0.017449 | FALSE |
| CECR2 | -1.811612183 | 0.163390509 | 0.762309376 | -2.376478948 | 0.017479 | FALSE |
| ZC2HC1C | 1.584299765 | 4.875875924 | 0.666683657 | 2.376389085 | 0.017483 | FALSE |
| IL12RB1 | 1.764752079 | 5.840124291 | 0.742794761 | 2.375827311 | 0.01751 | FALSE |
| EHBP1 | -1.355650239 | 0.257779621 | 0.571082145 | -2.373827042 | 0.017605 | TRUE |
| AHSA2 | 1.638247131 | 5.146141091 | 0.690207049 | 2.37355897 | 0.017618 | FALSE |
| SLC2A14 | 1.368529324 | 3.929567325 | 0.576934137 | 2.372072021 | 0.017689 | TRUE |
| PIWIL2 | -1.742640386 | 0.17505757 | 0.734840348 | -2.371454414 | 0.017718 | TRUE |

| | | | | | | |
|---|---|---|---|---|---|---|
| DOK2 | 0.837323848 | 2.310176315 | 0.353221371 | 2.37053564 | 0.017762 | TRUE |
| UBE2E2 | -1.162180254 | 0.312803446 | 0.490612138 | -2.368837142 | 0.017844 | TRUE |
| MAN1A2 | -1.317001096 | 0.267937618 | 0.557189159 | -2.363651688 | 0.018096 | TRUE |
| TRPV2 | 1.415205579 | 4.117332817 | 0.599299532 | 2.361432812 | 0.018204 | TRUE |
| MAPKAPK2 | 1.803914925 | 6.073377804 | 0.764323736 | 2.360145106 | 0.018268 | TRUE |
| SERTAD2 | -1.995887995 | 0.135892928 | 0.846209978 | -2.358620257 | 0.018343 | TRUE |
| HDHD3 | 1.230169929 | 3.421810952 | 0.521566379 | 2.358606647 | 0.018344 | TRUE |
| CAMK2G | 1.714669978 | 5.554841992 | 0.727932497 | 2.355534319 | 0.018496 | TRUE |
| RIC8B | -1.566388306 | 0.208797936 | 0.665164088 | -2.354890071 | 0.018528 | TRUE |
| RPL10A | -1.245927113 | 0.287674078 | 0.529399955 | -2.353470378 | 0.018599 | TRUE |
| TEX2 | -1.890367217 | 0.151016343 | 0.803332035 | -2.353158016 | 0.018615 | FALSE |
| FAM126B | -1.850244579 | 0.157198714 | 0.786327059 | -2.353021631 | 0.018622 | TRUE |
| TPRKB | -1.363182059 | 0.255845365 | 0.579582247 | -2.352007961 | 0.018672 | TRUE |
| AAMDC | -1.463335891 | 0.231462851 | 0.622307415 | -2.351467869 | 0.0187 | TRUE |
| PPIP5K2 | -1.347590093 | 0.259865759 | 0.573094117 | -2.351428942 | 0.018701 | TRUE |
| DROSHA | -1.438151721 | 0.237366072 | 0.612245644 | -2.348978284 | 0.018825 | FALSE |
| COQ3 | -1.39025163 | 0.249012638 | 0.592237438 | -2.347456512 | 0.018902 | FALSE |
| TANGO2 | 1.238180534 | 3.449331811 | 0.527582613 | 2.346894125 | 0.018931 | TRUE |
| CNN3 | -1.196997874 | 0.302099793 | 0.51012609 | -2.346474522 | 0.018952 | FALSE |
| C1orf115 | -1.559093763 | 0.210326591 | 0.664520636 | -2.34619315 | 0.018966 | FALSE |
| RUNX3 | 1.20227134 | 3.327666609 | 0.512468739 | 2.346038399 | 0.018974 | TRUE |
| RELT | 1.015872952 | 2.761773241 | 0.433018543 | 2.346026443 | 0.018975 | TRUE |
| CLDN16 | -1.254603683 | 0.285188851 | 0.534884487 | -2.345560048 | 0.018999 | FALSE |
| CORO1A | 0.931720179 | 2.538872739 | 0.397309296 | 2.345075203 | 0.019023 | TRUE |
| NMRK1 | -1.540483433 | 0.214277488 | 0.657560246 | -2.342725921 | 0.019143 | FALSE |
| DPP3 | 1.429237257 | 4.175513133 | 0.610287057 | 2.341909829 | 0.019185 | TRUE |
| METRNL | 0.804022221 | 2.234510574 | 0.343433314 | 2.341130548 | 0.019225 | TRUE |
| ZNF599 | -1.258516508 | 0.284075137 | 0.538268145 | -2.338084689 | 0.019383 | FALSE |
| RUFY4 | 1.432982944 | 4.191182628 | 0.61303853 | 2.337508777 | 0.019413 | FALSE |
| ICAM5 | 0.73781208 | 2.091354789 | 0.315765736 | 2.33658056 | 0.019461 | TRUE |
| FRS2 | -1.34398512 | 0.260804259 | 0.57529879 | -2.336151478 | 0.019483 | TRUE |
| IL10 | 1.455032081 | 4.284620919 | 0.622942092 | 2.335742119 | 0.019505 | TRUE |
| FAM96A | -1.39310808 | 0.24830236 | 0.596565509 | -2.335213917 | 0.019532 | FALSE |
| KLK12 | 1.32449616 | 3.760290294 | 0.56721692 | 2.33507872 | 0.019539 | TRUE |
| ZDHHC1 | 1.263147105 | 3.536533836 | 0.541183035 | 2.334047862 | 0.019593 | TRUE |
| ZRANB2 | -1.347919155 | 0.259780261 | 0.577658992 | -2.333416729 | 0.019626 | FALSE |
| EFCAB2 | -1.760019026 | 0.17204159 | 0.754532885 | -2.332594193 | 0.019669 | TRUE |
| EIF4E1B | 1.435075324 | 4.199961355 | 0.615544174 | 2.331392913 | 0.019733 | TRUE |
| CCDC12 | 1.778437416 | 5.920597754 | 0.762942863 | 2.331023073 | 0.019752 | TRUE |
| CWC22 | -1.317310119 | 0.267854832 | 0.565503131 | -2.329447966 | 0.019835 | FALSE |
| TNFSF18 | -1.384043602 | 0.250563323 | 0.594164973 | -2.329392786 | 0.019838 | FALSE |
| ATP6V0D2 | -1.400445949 | 0.246487019 | 0.60138531 | -2.328699963 | 0.019875 | FALSE |
| GMPPB | 1.201110248 | 3.323805123 | 0.51582435 | 2.328525684 | 0.019884 | TRUE |
| SLC25A33 | -1.78673497 | 0.167506191 | 0.767404714 | -2.328282506 | 0.019897 | FALSE |

| | | | | | | |
|---|---|---|---|---|---|---|
| UNC13D | 0.831167844 | 2.295998543 | 0.357058547 | 2.327819485 | 0.019922 | TRUE |
| CALHM1 | 1.169558288 | 3.220569759 | 0.502674532 | 2.326671064 | 0.019983 | TRUE |
| ADA | 1.595120947 | 4.928925173 | 0.685693149 | 2.326289755 | 0.020003 | TRUE |
| SPEF1 | 1.243204356 | 3.466704241 | 0.534935631 | 2.324026076 | 0.020124 | FALSE |
| SND1 | -2.13153317 | 0.118655236 | 0.917704702 | -2.322678707 | 0.020196 | TRUE |
| TMIGD2 | 1.336624698 | 3.806174811 | 0.576314162 | 2.319264016 | 0.020381 | TRUE |
| REV3L | -1.344254051 | 0.26073413 | 0.579964099 | -2.317822869 | 0.020459 | TRUE |
| RSL24D1 | -1.311346025 | 0.269457116 | 0.566025883 | -2.316759823 | 0.020517 | TRUE |
| FAT4 | -1.171106711 | 0.310023645 | 0.505611678 | -2.316217689 | 0.020546 | TRUE |
| SLC34A3 | 0.849729925 | 2.339015056 | 0.366923838 | 2.315820988 | 0.020568 | TRUE |
| PRPH2 | 1.501644569 | 4.489065582 | 0.648782872 | 2.314556433 | 0.020637 | FALSE |
| SYT6 | 1.471313435 | 4.354951334 | 0.635690487 | 2.314512275 | 0.02064 | TRUE |
| ASB4 | -1.269161161 | 0.281067293 | 0.548364827 | -2.314446694 | 0.020643 | TRUE |
| HIBCH | -1.384749437 | 0.250386529 | 0.598538629 | -2.313550654 | 0.020692 | FALSE |
| GPR137C | -1.284792621 | 0.276707961 | 0.555964711 | -2.310924768 | 0.020837 | FALSE |
| MALSU1 | -1.433567306 | 0.238456755 | 0.620513449 | -2.310292081 | 0.020872 | TRUE |
| ASB1 | 1.445230637 | 4.242830583 | 0.626080068 | 2.308379889 | 0.020978 | FALSE |
| ENSA | 1.579509856 | 4.852576767 | 0.684697583 | 2.306872253 | 0.021062 | TRUE |
| IMMP2L | -1.006980768 | 0.365320303 | 0.437053144 | -2.304023622 | 0.021221 | FALSE |
| PGM3 | -1.51529839 | 0.219742606 | 0.657702108 | -2.303928132 | 0.021227 | FALSE |
| MLLT6 | 1.128404825 | 3.090722323 | 0.489836513 | 2.303635591 | 0.021243 | TRUE |
| FZD5 | 0.935764613 | 2.549161835 | 0.406244617 | 2.303451106 | 0.021253 | TRUE |
| MED29 | 1.298970839 | 3.665522315 | 0.564388546 | 2.301554221 | 0.02136 | TRUE |
| PLS1 | -1.494198011 | 0.224428522 | 0.649771113 | -2.29957593 | 0.021472 | FALSE |
| DOK1 | 1.077482232 | 2.937274857 | 0.468907589 | 2.297856248 | 0.02157 | FALSE |
| KLRF2 | -1.368215728 | 0.25456076 | 0.595624785 | -2.297110132 | 0.021612 | FALSE |
| ASIC3 | -1.196198296 | 0.302341442 | 0.520951604 | -2.296179311 | 0.021666 | FALSE |
| PCSK7 | 1.519460093 | 4.569757287 | 0.661947246 | 2.295439859 | 0.021708 | FALSE |
| PTTG1IP | 1.301431849 | 3.674554308 | 0.567262253 | 2.294233119 | 0.021777 | TRUE |
| C11orf85 | 1.707272943 | 5.513904227 | 0.744496715 | 2.293190699 | 0.021837 | FALSE |
| RRAS2 | -1.086217932 | 0.337490495 | 0.474051105 | -2.291351969 | 0.021943 | TRUE |
| COIL | -1.568635104 | 0.208329336 | 0.685219809 | -2.289243659 | 0.022065 | FALSE |
| CREB3L1 | 1.308135996 | 3.699271824 | 0.571541922 | 2.288783981 | 0.022092 | TRUE |
| ORMDL1 | -1.22414114 | 0.294010106 | 0.535252386 | -2.287035372 | 0.022194 | TRUE |
| NIT2 | -1.429278151 | 0.23948173 | 0.625295074 | -2.285765888 | 0.022268 | TRUE |
| ENAH | -1.418552574 | 0.242064133 | 0.621088318 | -2.283978838 | 0.022373 | FALSE |
| NMNAT2 | -1.196480502 | 0.302256132 | 0.523990712 | -2.283400211 | 0.022407 | TRUE |
| ABO | 0.819522916 | 2.269416876 | 0.358955411 | 2.2830772 | 0.022426 | TRUE |
| TLCD1 | 1.369839971 | 3.934720976 | 0.60016869 | 2.282424914 | 0.022464 | FALSE |
| PGPEP1L | -1.386201031 | 0.250023334 | 0.607411177 | -2.282146073 | 0.022481 | FALSE |
| LIMCH1 | -1.584208219 | 0.205110131 | 0.694470583 | -2.281173974 | 0.022538 | TRUE |
| SLC38A2 | -1.153322699 | 0.315586427 | 0.506268215 | -2.278086326 | 0.022721 | FALSE |
| THUMPD3 | -1.256539569 | 0.284637292 | 0.552245825 | -2.275326515 | 0.022886 | TRUE |
| TRIM54 | 1.491551762 | 4.443986182 | 0.655729289 | 2.27464563 | 0.022927 | TRUE |

| | | | | | | |
|---|---|---|---|---|---|---|
| FBXO11 | -1.536442228 | 0.215145179 | 0.675733333 | -2.273740473 | 0.022982 | FALSE |
| HYAL2 | 1.231984521 | 3.428025778 | 0.542123782 | 2.272515175 | 0.023055 | FALSE |
| MBD3L1 | -1.302712998 | 0.271793417 | 0.573404738 | -2.271890885 | 0.023093 | TRUE |
| RARS | -1.323122007 | 0.266302604 | 0.58251584 | -2.271392324 | 0.023123 | FALSE |
| TMTC1 | -0.798615252 | 0.449951602 | 0.352078069 | -2.268290255 | 0.023312 | TRUE |
| DNA2 | -1.57020492 | 0.208002554 | 0.692346761 | -2.267945786 | 0.023333 | FALSE |
| ATAD3A | 1.067739274 | 2.908796071 | 0.470798312 | 2.267933521 | 0.023333 | TRUE |
| PRR22 | 1.09231409 | 2.981164781 | 0.481783275 | 2.267231235 | 0.023376 | FALSE |
| MNAT1 | -1.465677516 | 0.230921485 | 0.647109455 | -2.264960748 | 0.023515 | TRUE |
| LRP8 | 1.200161925 | 3.320654576 | 0.530425414 | 2.262640314 | 0.023658 | TRUE |
| UBR5 | -1.57331135 | 0.207357411 | 0.695864268 | -2.260945737 | 0.023763 | FALSE |
| GINM1 | -1.557411598 | 0.210680692 | 0.688892182 | -2.260747965 | 0.023775 | TRUE |
| GPRC5C | 1.078156899 | 2.939257206 | 0.477083012 | 2.25989371 | 0.023828 | TRUE |
| RPL31 | 1.425185973 | 4.158631165 | 0.630750203 | 2.259509339 | 0.023852 | FALSE |
| GCC1 | 1.495258424 | 4.460489099 | 0.662098781 | 2.25836154 | 0.023923 | TRUE |
| CXCR4 | 1.231034384 | 3.424770232 | 0.545133052 | 2.25822738 | 0.023931 | TRUE |
| RGS19 | 0.817258743 | 2.264284337 | 0.361970221 | 2.257806568 | 0.023958 | TRUE |
| IPO7 | -1.919780462 | 0.146639151 | 0.850458385 | -2.257347915 | 0.023986 | FALSE |
| SLC4A7 | -1.45842535 | 0.232602254 | 0.64658925 | -2.25556696 | 0.024098 | TRUE |
| IFT88 | -1.550949274 | 0.212046588 | 0.687730662 | -2.255169591 | 0.024123 | TRUE |
| FGF2 | -1.095512271 | 0.334368276 | 0.485801341 | -2.255062264 | 0.024129 | FALSE |
| INADL | -1.820584705 | 0.161931041 | 0.807623322 | -2.25424979 | 0.02418 | FALSE |
| SEC24B | -1.322141507 | 0.266563842 | 0.587165339 | -2.251736299 | 0.024339 | TRUE |
| MPDZ | -1.041352285 | 0.352977033 | 0.462520524 | -2.251472596 | 0.024356 | TRUE |
| WDR88 | 1.333245782 | 3.793335767 | 0.592454209 | 2.25037777 | 0.024425 | FALSE |
| C12orf54 | 1.327182202 | 3.770404167 | 0.59018941 | 2.248739436 | 0.024529 | TRUE |
| ARL6IP6 | -1.225952038 | 0.293478165 | 0.545238723 | -2.248468398 | 0.024546 | FALSE |
| HEXIM2 | 1.001018552 | 2.72105195 | 0.44538978 | 2.247511275 | 0.024607 | TRUE |
| ACSM5 | -1.060087553 | 0.346425478 | 0.471694061 | -2.247404918 | 0.024614 | FALSE |
| RFXAP | -1.24229061 | 0.28872211 | 0.552769802 | -2.247392323 | 0.024615 | FALSE |
| MROH6 | 0.693174313 | 2.000054266 | 0.308536084 | 2.246655576 | 0.024662 | TRUE |
| ITGAE | 1.545834094 | 4.691883466 | 0.688091985 | 2.246551517 | 0.024669 | TRUE |
| HRC | 1.201525823 | 3.325186699 | 0.534978904 | 2.245931222 | 0.024708 | FALSE |
| RYR1 | 1.249498476 | 3.488592905 | 0.556393552 | 2.245709841 | 0.024723 | TRUE |
| PHLDB2 | -1.200711326 | 0.300980041 | 0.534855327 | -2.2449273 | 0.024773 | FALSE |
| SNX4 | -1.581769364 | 0.205610976 | 0.704909768 | -2.243931685 | 0.024837 | FALSE |
| TMUB1 | 0.894196124 | 2.445369225 | 0.398549576 | 2.243625831 | 0.024856 | TRUE |
| UBN2 | -1.413510598 | 0.243287697 | 0.630350137 | -2.24242134 | 0.024934 | TRUE |
| MFSD10 | 0.787398199 | 2.19767108 | 0.351152084 | 2.242328141 | 0.02494 | FALSE |
| ZNF844 | -1.323640546 | 0.266164552 | 0.590486283 | -2.241610997 | 0.024987 | FALSE |
| ACO1 | -1.353563068 | 0.258318214 | 0.603849106 | -2.241558452 | 0.02499 | FALSE |
| ARHGEF2 | 1.541358458 | 4.670931229 | 0.687957693 | 2.240484369 | 0.025059 | TRUE |
| AGO3 | -1.52401367 | 0.217835809 | 0.680401754 | -2.239873221 | 0.025099 | TRUE |
| NR1H3 | 1.412567403 | 4.106484884 | 0.630678222 | 2.239759284 | 0.025107 | TRUE |

| | | | | | | |
|---|---|---|---|---|---|---|
| PTPRJ | 1.949976298 | 7.028520988 | 0.870637942 | 2.239709763 | 0.02511 | TRUE |
| ZNF626 | -1.297872975 | 0.273112092 | 0.579484927 | -2.239701008 | 0.02511 | TRUE |
| PUS7L | -1.42772527 | 0.239853905 | 0.637939404 | -2.238026465 | 0.025219 | FALSE |
| PKP2 | -1.329502959 | 0.26460875 | 0.594733941 | -2.235458359 | 0.025387 | TRUE |
| CETN3 | -1.17704125 | 0.308189246 | 0.52726403 | -2.232356432 | 0.025591 | FALSE |
| CEP57 | -1.482747394 | 0.227013136 | 0.664432783 | -2.23159879 | 0.025641 | TRUE |
| ARHGAP32 | -1.079296337 | 0.33983457 | 0.483789637 | -2.230920747 | 0.025686 | FALSE |
| CTD-2510F5.6 | 1.469127052 | 4.345440143 | 0.658682727 | 2.230401668 | 0.025721 | FALSE |
| PUS3 | 1.329954462 | 3.780871209 | 0.596396348 | 2.22998425 | 0.025748 | FALSE |
| SERPINF2 | 1.075471211 | 2.93137387 | 0.482521063 | 2.228858577 | 0.025823 | TRUE |
| EMILIN1 | 0.66792505 | 1.95018658 | 0.299828097 | 2.227693327 | 0.025901 | TRUE |
| KAT6B | -1.909333069 | 0.148179179 | 0.857615287 | -2.226328165 | 0.025992 | FALSE |
| KIF5B | -1.076418317 | 0.34081403 | 0.483695078 | -2.225406802 | 0.026054 | TRUE |
| CA3 | -1.273756491 | 0.279778659 | 0.572632304 | -2.224388115 | 0.026122 | TRUE |
| PACRGL | -1.402908877 | 0.245880686 | 0.630955429 | -2.22346748 | 0.026184 | TRUE |
| SBF2 | -1.534676259 | 0.215525454 | 0.690933306 | -2.221164107 | 0.02634 | TRUE |
| PPARG | -1.091157364 | 0.335827594 | 0.491465549 | -2.220211297 | 0.026404 | TRUE |
| DOCK1 | -1.652454784 | 0.191579046 | 0.744406132 | -2.219829625 | 0.02643 | FALSE |
| PNPLA6 | 0.872390576 | 2.392623772 | 0.39311989 | 2.219146367 | 0.026477 | TRUE |
| ADRB2 | 1.203715182 | 3.332474704 | 0.542479941 | 2.218911873 | 0.026493 | FALSE |
| INO80 | -1.934730436 | 0.144463206 | 0.872155008 | -2.218333231 | 0.026532 | TRUE |
| PIK3C3 | -1.019679027 | 0.3607107 | 0.459878206 | -2.217280604 | 0.026604 | FALSE |
| RASA2 | -1.253363612 | 0.285542725 | 0.565738951 | -2.215445145 | 0.02673 | FALSE |
| PPP3CB | -1.606766974 | 0.200534902 | 0.72537827 | -2.215074589 | 0.026755 | FALSE |
| ZNF812 | 1.326470408 | 3.767721372 | 0.599274262 | 2.213461337 | 0.026866 | FALSE |
| TTC23L | 1.375613046 | 3.95750211 | 0.622055738 | 2.21139837 | 0.027008 | FALSE |
| DYRK4 | 1.433029625 | 4.19137828 | 0.648069389 | 2.211228689 | 0.02702 | TRUE |
| EMILIN2 | 1.666264926 | 5.292363465 | 0.754102068 | 2.209601323 | 0.027133 | FALSE |
| ARID4B | -1.698526553 | 0.182952897 | 0.768831519 | -2.209231165 | 0.027159 | FALSE |
| PPFIBP2 | -1.019714626 | 0.360697859 | 0.46171217 | -2.208550461 | 0.027206 | TRUE |
| ACTB | 1.192441437 | 3.295116215 | 0.540508662 | 2.206146766 | 0.027374 | TRUE |
| SLC8B1 | 1.615896327 | 5.032396575 | 0.732617222 | 2.205648841 | 0.027409 | TRUE |
| DCUN1D2 | -1.673443442 | 0.187599963 | 0.758907172 | -2.205070006 | 0.027449 | FALSE |
| SFSWAP | 1.889130352 | 6.613614666 | 0.856752929 | 2.204988497 | 0.027455 | TRUE |
| AMT | 1.203428285 | 3.331518763 | 0.545849944 | 2.204687017 | 0.027476 | TRUE |
| MYO9A | -1.45305836 | 0.233853984 | 0.659551776 | -2.20309976 | 0.027588 | FALSE |
| LEPR | -1.310776113 | 0.269610727 | 0.595141894 | -2.202459828 | 0.027633 | TRUE |
| PRSS36 | 0.913009382 | 2.491810069 | 0.414640985 | 2.201927485 | 0.02767 | TRUE |
| FAM221A | -1.384349188 | 0.250486766 | 0.628855322 | -2.201379459 | 0.027709 | TRUE |
| HIGD2B | -1.345424822 | 0.260429048 | 0.61131972 | -2.200852969 | 0.027746 | TRUE |
| EXOC6B | -1.278689513 | 0.278401904 | 0.581152491 | -2.200265045 | 0.027788 | TRUE |
| SNIP1 | 1.282127676 | 3.604300356 | 0.582752818 | 2.200122655 | 0.027798 | FALSE |
| DHX35 | -1.617095388 | 0.198474354 | 0.735143644 | -2.199699883 | 0.027828 | TRUE |
| PHYHIPL | -0.985219006 | 0.373357451 | 0.448028396 | -2.199010183 | 0.027877 | FALSE |

| | | | | | | |
|---|---|---|---|---|---|---|
| MAP3K13 | -1.469268761 | 0.230093677 | 0.668366378 | -2.198298432 | 0.027928 | TRUE |
| SMNDC1 | -1.224923186 | 0.293780266 | 0.557503456 | -2.197158011 | 0.028009 | FALSE |
| CRNN | -1.099970213 | 0.332880999 | 0.50068757 | -2.196919356 | 0.028026 | TRUE |
| ORC4 | -1.182023716 | 0.306657522 | 0.538089057 | -2.196706475 | 0.028041 | FALSE |
| DIS3 | -1.23655338 | 0.290383336 | 0.563168314 | -2.19570837 | 0.028113 | TRUE |
| MFSD12 | 0.988352151 | 2.686803377 | 0.450252118 | 2.195108277 | 0.028156 | TRUE |
| ATP4B | 1.233212736 | 3.43223872 | 0.5619823 | 2.194397823 | 0.028207 | TRUE |
| DNAJC2 | -1.419245792 | 0.241896388 | 0.646887605 | -2.193960405 | 0.028238 | TRUE |
| RAPGEF4 | -1.231655269 | 0.291809155 | 0.561522514 | -2.193420992 | 0.028277 | TRUE |
| TRMT61A | 0.907653889 | 2.478500868 | 0.413854118 | 2.193173508 | 0.028295 | FALSE |
| CHMP4B | 1.483496273 | 4.4083315 | 0.676529113 | 2.192804779 | 0.028321 | FALSE |
| BIRC2 | -1.343756378 | 0.260863922 | 0.6129308 | -2.192345983 | 0.028355 | FALSE |
| XDH | -1.181428587 | 0.306840077 | 0.539027415 | -2.191778293 | 0.028396 | FALSE |
| TRIM45 | -1.374952037 | 0.252851723 | 0.627450941 | -2.191329948 | 0.028428 | FALSE |
| CEPT1 | -1.134615863 | 0.321545615 | 0.517853893 | -2.190996107 | 0.028452 | TRUE |
| ATP5C1 | -1.359277226 | 0.256846352 | 0.620854316 | -2.189365834 | 0.02857 | TRUE |
| ADGRV1 | -1.323942096 | 0.266084302 | 0.604842414 | -2.188904192 | 0.028604 | TRUE |
| KIF17 | 1.223324024 | 3.398465561 | 0.558967552 | 2.188542107 | 0.02863 | TRUE |
| EPOR | 1.102555755 | 3.011853757 | 0.503852 | 2.188253208 | 0.028651 | TRUE |
| PRDM13 | 1.152224451 | 3.165225975 | 0.526999027 | 2.186388193 | 0.028787 | FALSE |
| ZBTB42 | 1.015049138 | 2.75949899 | 0.464558378 | 2.1849765 | 0.028891 | FALSE |
| TMC8 | 0.80657013 | 2.240211161 | 0.369497766 | 2.182882292 | 0.029044 | TRUE |
| BCL2L1 | 1.387117364 | 4.003293367 | 0.635464929 | 2.182838582 | 0.029048 | TRUE |
| CCDC172 | -1.164533853 | 0.312068098 | 0.533531655 | -2.182689333 | 0.029059 | FALSE |
| LRRC28 | -1.379188221 | 0.251782862 | 0.632189078 | -2.181607162 | 0.029139 | TRUE |
| FARP1 | -1.03031707 | 0.356893782 | 0.47237082 | -2.18116155 | 0.029171 | TRUE |
| SACM1L | -1.130666192 | 0.322818126 | 0.518436413 | -2.180915852 | 0.02919 | TRUE |
| DIRAS2 | -1.264368128 | 0.282417692 | 0.579775692 | -2.180788441 | 0.029199 | FALSE |
| ARRB1 | 1.376343513 | 3.960393991 | 0.631157842 | 2.180664519 | 0.029208 | FALSE |
| PTAR1 | -1.093047306 | 0.335193499 | 0.501462073 | -2.179720792 | 0.029278 | TRUE |
| CDK20 | -1.241707898 | 0.288890401 | 0.569988527 | -2.178478758 | 0.02937 | TRUE |
| MSH3 | -1.387689152 | 0.249651545 | 0.637380939 | -2.17717391 | 0.029468 | TRUE |
| NEFH | 1.205156154 | 3.337280168 | 0.553675674 | 2.176646385 | 0.029507 | FALSE |
| UBE4A | -1.368324021 | 0.254533194 | 0.628714229 | -2.176384688 | 0.029527 | TRUE |
| TUBE1 | -1.231404204 | 0.291882427 | 0.565804212 | -2.176378643 | 0.029527 | FALSE |
| EHD1 | 0.956508217 | 2.6025929 | 0.439536418 | 2.176175119 | 0.029542 | TRUE |
| C14orf159 | 2.160642888 | 8.676714024 | 0.993526633 | 2.174720654 | 0.029651 | TRUE |
| LIG1 | 1.24967211 | 3.489198698 | 0.575130223 | 2.172850704 | 0.029792 | FALSE |
| CYP11A1 | 1.32492946 | 3.76191998 | 0.609834285 | 2.172605728 | 0.02981 | FALSE |
| ORC3 | -1.518627628 | 0.219012247 | 0.69933153 | -2.171541771 | 0.02989 | FALSE |
| OPRL1 | 0.885851238 | 2.425047807 | 0.407980429 | 2.171308167 | 0.029908 | FALSE |
| CYP2D6 | 0.983751048 | 2.674469513 | 0.45341711 | 2.169638124 | 0.030034 | TRUE |
| RPAP2 | -1.365434146 | 0.255269827 | 0.62955704 | -2.168880753 | 0.030092 | TRUE |
| PDE4DIP | -1.296786383 | 0.273409015 | 0.598048348 | -2.168363791 | 0.030131 | FALSE |

| | | | | | | |
|---|---|---|---|---|---|---|
| SH2D4A | -1.559287147 | 0.210285921 | 0.719192534 | -2.168108083 | 0.03015 | FALSE |
| CTCFL | 1.534965386 | 4.641164876 | 0.708062939 | 2.167837493 | 0.030171 | TRUE |
| GRIK1 | -1.268679922 | 0.281202586 | 0.585770581 | -2.165830723 | 0.030324 | FALSE |
| TK2 | 1.613299371 | 5.019344618 | 0.744978972 | 2.165563634 | 0.030345 | TRUE |
| TMEM260 | -1.51544165 | 0.219711128 | 0.700358093 | -2.16380972 | 0.030479 | TRUE |
| USP53 | -1.211877302 | 0.297637998 | 0.560094063 | -2.163703172 | 0.030487 | TRUE |
| DEDD | -1.451184603 | 0.23429258 | 0.67081372 | -2.163319799 | 0.030517 | TRUE |
| OGFOD3 | 1.078246 | 2.939519109 | 0.498482095 | 2.163058634 | 0.030537 | TRUE |
| CYB5D2 | 1.211020584 | 3.356908911 | 0.560147405 | 2.161967675 | 0.030621 | TRUE |
| ASPDH | 0.992199265 | 2.697159723 | 0.4590596 | 2.161373521 | 0.030666 | FALSE |
| PPP1R1B | 0.896432452 | 2.450843993 | 0.414759418 | 2.161331156 | 0.03067 | TRUE |
| ARHGAP5 | -1.234683274 | 0.290926892 | 0.571375223 | -2.160897472 | 0.030703 | TRUE |
| ST6GALNAC4 | 0.999956814 | 2.71816444 | 0.462809203 | 2.160624308 | 0.030724 | TRUE |
| ADGRA3 | -1.221277614 | 0.294853218 | 0.565483981 | -2.159703288 | 0.030796 | TRUE |
| C9orf85 | -1.422728659 | 0.241055361 | 0.659156802 | -2.158407005 | 0.030896 | FALSE |
| TCP1 | -1.295970725 | 0.273632114 | 0.600500764 | -2.158150002 | 0.030916 | FALSE |
| TSPO | 0.946841336 | 2.577555157 | 0.438778548 | 2.157902524 | 0.030935 | FALSE |
| PPP2R1B | -1.504894741 | 0.222040664 | 0.697503028 | -2.157545818 | 0.030963 | TRUE |
| STX18 | -1.671722871 | 0.187923019 | 0.775409937 | -2.155921392 | 0.03109 | TRUE |
| ZNF814 | -1.172352287 | 0.309637727 | 0.5443753 | -2.153573624 | 0.031274 | TRUE |
| ABCB4 | -1.161841025 | 0.312909576 | 0.539524196 | -2.153454902 | 0.031283 | TRUE |
| PRMT3 | -1.062229742 | 0.345684164 | 0.493462172 | -2.152606221 | 0.03135 | FALSE |
| NEDD4L | -1.614429895 | 0.199004091 | 0.750143289 | -2.152162019 | 0.031385 | TRUE |
| CNTLN | -1.177763752 | 0.307966659 | 0.547295094 | -2.151972063 | 0.0314 | FALSE |
| YIF1B | 0.95871019 | 2.608330052 | 0.445540822 | 2.151789784 | 0.031414 | TRUE |
| CPB2 | -1.214833217 | 0.296759505 | 0.564590522 | -2.151706713 | 0.03142 | TRUE |
| COX5B | 1.18843936 | 3.281955259 | 0.552339832 | 2.151645222 | 0.031425 | TRUE |
| P2RX6 | 1.176540887 | 3.243136402 | 0.546868308 | 2.151415376 | 0.031443 | TRUE |
| THOC7 | -1.238445953 | 0.289834284 | 0.575775812 | -2.15091695 | 0.031483 | TRUE |
| TAF2 | -1.38798444 | 0.249577837 | 0.645422714 | -2.150504483 | 0.031515 | FALSE |
| GSTO2 | -1.293891874 | 0.274201546 | 0.602174877 | -2.148697866 | 0.031658 | FALSE |
| STARD3 | 0.782519666 | 2.186975779 | 0.364218542 | 2.148489374 | 0.031675 | TRUE |
| DHRS13 | 1.353828821 | 3.872223235 | 0.630169252 | 2.148357472 | 0.031685 | TRUE |
| GGA3 | 1.320700095 | 3.746043046 | 0.614803646 | 2.148165685 | 0.031701 | FALSE |
| PYCRL | 0.999341736 | 2.71649307 | 0.465226813 | 2.148074246 | 0.031708 | TRUE |
| MIIP | 1.118427624 | 3.060038888 | 0.520854606 | 2.147293334 | 0.03177 | TRUE |
| TGFB1 | 1.112276366 | 3.041273572 | 0.518620546 | 2.144682417 | 0.031978 | TRUE |
| CUEDC1 | 1.607289535 | 4.989269645 | 0.749669066 | 2.143998742 | 0.032033 | FALSE |
| CAMKK2 | 1.516352251 | 4.555577249 | 0.707487976 | 2.143290491 | 0.03209 | FALSE |
| TRIQK | -1.034282562 | 0.355481325 | 0.482931988 | -2.141673337 | 0.03222 | TRUE |
| HOOK1 | -1.079233263 | 0.339856006 | 0.503990277 | -2.14137715 | 0.032244 | TRUE |
| PITPNM3 | 1.26451725 | 3.541382721 | 0.590700165 | 2.140709152 | 0.032297 | FALSE |
| SRPK1 | -1.235328618 | 0.290739205 | 0.57708067 | -2.140651528 | 0.032302 | FALSE |
| LHX4 | 1.326621206 | 3.768289579 | 0.619801686 | 2.140396253 | 0.032323 | FALSE |

| | | | | | | |
|---|---|---|---|---|---|---|
| IKBKE | 1.457040529 | 4.293235004 | 0.68110551 | 2.139228808 | 0.032417 | FALSE |
| HIBADH | -1.289331491 | 0.275454866 | 0.602895202 | -2.138566516 | 0.032471 | TRUE |
| ADCY4 | 1.134783223 | 3.110499186 | 0.530753776 | 2.138059631 | 0.032512 | TRUE |
| MGMT | -1.162512872 | 0.312699419 | 0.543735637 | -2.138011182 | 0.032516 | FALSE |
| RPL18A | 1.242416734 | 3.463974863 | 0.581316664 | 2.137246032 | 0.032578 | FALSE |
| CPS1 | -0.730473563 | 0.48168083 | 0.341806154 | -2.13709892 | 0.03259 | FALSE |
| ZBTB38 | -1.770609287 | 0.170229239 | 0.828537501 | -2.137029748 | 0.032596 | TRUE |
| TFCP2 | -1.673564205 | 0.187577309 | 0.783881968 | -2.134969643 | 0.032763 | FALSE |
| OTUD4 | -1.237733727 | 0.290040785 | 0.579761623 | -2.1349011 | 0.032769 | TRUE |
| ZHX3 | -1.584051372 | 0.205142305 | 0.742103681 | -2.13454186 | 0.032798 | FALSE |
| UBXN6 | 0.73169279 | 2.078596258 | 0.342908037 | 2.133787229 | 0.03286 | TRUE |
| DPH6 | -1.050790136 | 0.34966136 | 0.49272308 | -2.132618052 | 0.032956 | TRUE |
| CCDC18 | -1.363355128 | 0.25580109 | 0.639795412 | -2.130923575 | 0.033095 | FALSE |
| MFSD2B | 0.876375137 | 2.402176345 | 0.411297606 | 2.130756717 | 0.033109 | TRUE |
| PRKCD | 1.04284501 | 2.837277626 | 0.489474611 | 2.130539534 | 0.033127 | TRUE |
| SLC2A1 | 1.329649585 | 3.779718686 | 0.624184833 | 2.130217711 | 0.033154 | TRUE |
| FAM46B | 1.127796728 | 3.088843436 | 0.529490492 | 2.129965966 | 0.033174 | TRUE |
| CEP95 | -1.342731602 | 0.261131386 | 0.630571857 | -2.129387137 | 0.033222 | TRUE |
| NXPE1 | -1.161655035 | 0.312967779 | 0.545582374 | -2.129201915 | 0.033238 | FALSE |
| PHOX2A | 1.094892057 | 2.988860039 | 0.51439892 | 2.128488248 | 0.033297 | TRUE |
| SFRP5 | 1.256442319 | 3.512901446 | 0.590947714 | 2.126148031 | 0.033491 | FALSE |
| HKDC1 | 1.381582981 | 3.981198804 | 0.650036716 | 2.125392222 | 0.033554 | FALSE |
| LRRC39 | -1.260608926 | 0.283481355 | 0.593553634 | -2.123833219 | 0.033684 | FALSE |
| STARD5 | -1.329392756 | 0.264637912 | 0.626194765 | -2.122970087 | 0.033756 | TRUE |
| ZBTB7B | 0.89112242 | 2.437864424 | 0.419837137 | 2.122543103 | 0.033792 | TRUE |
| RXRG | -1.38255246 | 0.250937228 | 0.651474625 | -2.122189273 | 0.033822 | TRUE |
| SMIM21 | -0.9911201 | 0.371160721 | 0.467030779 | -2.122172981 | 0.033823 | TRUE |
| MDN1 | -1.657575061 | 0.190600615 | 0.781109285 | -2.122078298 | 0.033831 | TRUE |
| GOLT1B | -1.189008496 | 0.30452305 | 0.560327677 | -2.121987802 | 0.033839 | TRUE |
| NT5DC2 | 1.023019358 | 2.781580686 | 0.482106419 | 2.121978297 | 0.03384 | FALSE |
| NEURL1 | 1.397537474 | 4.045226218 | 0.658848262 | 2.121182608 | 0.033906 | FALSE |
| KLK2 | 1.100546005 | 3.005806763 | 0.519026642 | 2.120403688 | 0.033972 | TRUE |
| CENPF | -1.32214412 | 0.266563145 | 0.623535509 | -2.120399082 | 0.033972 | FALSE |
| CPEB2 | -0.923747013 | 0.397028579 | 0.435684847 | -2.120218364 | 0.033988 | FALSE |
| CST7 | 0.94129058 | 2.563287413 | 0.443994912 | 2.120048124 | 0.034002 | TRUE |
| ARHGDIA | 0.678653728 | 1.971222145 | 0.320149858 | 2.119800183 | 0.034023 | TRUE |
| CFAP43 | -1.343175142 | 0.26101559 | 0.63371689 | -2.119519243 | 0.034047 | TRUE |
| PCOLCE2 | -1.297407894 | 0.273239141 | 0.612321702 | -2.118833759 | 0.034105 | TRUE |
| AGPAT2 | 0.6907783 | 1.995267847 | 0.326224769 | 2.117491886 | 0.034218 | TRUE |
| SLC22A6 | -1.260679628 | 0.283461313 | 0.595688914 | -2.116338913 | 0.034316 | TRUE |
| SLC16A5 | 1.234473339 | 3.436568138 | 0.583328501 | 2.116257542 | 0.034323 | FALSE |
| C12orf43 | -1.22685465 | 0.293213387 | 0.579890503 | -2.115666049 | 0.034373 | FALSE |
| NDST1 | 1.510206574 | 4.527665995 | 0.714664666 | 2.1131681 | 0.034586 | TRUE |
| CRISPLD2 | 1.376554714 | 3.961230518 | 0.65175116 | 2.112086328 | 0.034679 | TRUE |

| | | | | | | |
|---|---|---|---|---|---|---|
| PIGX | 1.41633301 | 4.121977442 | 0.670836246 | 2.1112947 | 0.034747 | FALSE |
| TRPM4 | 1.170787708 | 3.224531628 | 0.554555998 | 2.111216384 | 0.034754 | TRUE |
| C22orf15 | 1.033848775 | 2.81186728 | 0.489989353 | 2.109941308 | 0.034863 | TRUE |
| PTGES | 1.12335484 | 3.075153565 | 0.532490459 | 2.109624353 | 0.034891 | FALSE |
| HVCN1 | 1.27021629 | 3.561622824 | 0.602176903 | 2.109373979 | 0.034912 | FALSE |
| WNT10A | 1.130999103 | 3.098750927 | 0.536487855 | 2.108154161 | 0.035018 | TRUE |
| GBX1 | 1.203013025 | 3.330135603 | 0.570791416 | 2.107622838 | 0.035064 | TRUE |
| CDIP1 | 1.146710248 | 3.147820309 | 0.544121234 | 2.107453591 | 0.035078 | TRUE |
| SLC2A6 | 0.784346563 | 2.190974809 | 0.372471667 | 2.105788529 | 0.035223 | FALSE |
| FAM76B | -1.19084706 | 0.30396368 | 0.565706008 | -2.105063481 | 0.035286 | TRUE |
| BTF3L4 | -1.397240623 | 0.247278358 | 0.663777484 | -2.104983459 | 0.035293 | TRUE |
| HRK | 1.187797035 | 3.279847854 | 0.564730336 | 2.103299502 | 0.03544 | TRUE |
| GLYATL2 | -1.212040207 | 0.297589516 | 0.576690474 | -2.101717058 | 0.035578 | TRUE |
| CCDC88C | 1.273080387 | 3.571838279 | 0.605788609 | 2.101525793 | 0.035595 | FALSE |
| KCTD1 | -1.763194397 | 0.171496161 | 0.839085284 | -2.101329188 | 0.035612 | FALSE |
| MCEMP1 | 1.133786849 | 3.107401508 | 0.539559612 | 2.101318972 | 0.035613 | FALSE |
| CCDC77 | -1.550638897 | 0.212112412 | 0.737990087 | -2.101164941 | 0.035626 | FALSE |
| IPO8 | -1.16658615 | 0.311428298 | 0.555347839 | -2.100640477 | 0.035673 | TRUE |
| SPINT2 | 1.057084896 | 2.877969171 | 0.503223109 | 2.100628683 | 0.035674 | TRUE |
| LCOR | -1.428399248 | 0.239692303 | 0.680548756 | -2.098893334 | 0.035826 | TRUE |
| DHX8 | -1.623067805 | 0.197292515 | 0.773373053 | -2.098686784 | 0.035845 | FALSE |
| COMMD8 | -1.301186955 | 0.272208502 | 0.620121522 | -2.098277367 | 0.035881 | TRUE |
| SPINK2 | 1.314380191 | 3.722443066 | 0.626590275 | 2.097670909 | 0.035934 | TRUE |
| GSTK1 | 1.659718329 | 5.257829656 | 0.79146722 | 2.097014617 | 0.035992 | TRUE |
| ITGAV | -1.21852034 | 0.29566733 | 0.581156857 | -2.09671507 | 0.036019 | FALSE |
| CDK12 | 0.946007312 | 2.575406311 | 0.451233399 | 2.096492224 | 0.036039 | TRUE |
| C21orf2 | 0.756879453 | 2.131614028 | 0.361241267 | 2.095218685 | 0.036152 | TRUE |
| CAPRIN2 | -1.244318464 | 0.288137217 | 0.594100287 | -2.094458615 | 0.036219 | TRUE |
| FGGY | -1.271088835 | 0.280526009 | 0.607097705 | -2.093713786 | 0.036285 | TRUE |
| PCSK4 | 0.658451266 | 1.931798175 | 0.314508419 | 2.093588681 | 0.036297 | TRUE |
| COX20 | -1.189018581 | 0.304519979 | 0.567987237 | -2.093389611 | 0.036314 | FALSE |
| SGMS2 | -1.023385843 | 0.359376087 | 0.488910885 | -2.093195047 | 0.036332 | TRUE |
| ATG2B | -1.38735571 | 0.249734803 | 0.662884056 | -2.092908553 | 0.036357 | FALSE |
| SLC12A5 | 0.946901693 | 2.577710735 | 0.452580574 | 2.092227874 | 0.036418 | FALSE |
| CCDC158 | -1.320007845 | 0.267133206 | 0.630948345 | -2.092101286 | 0.036429 | FALSE |
| GOLGB1 | -1.251534131 | 0.286065598 | 0.599131289 | -2.088914657 | 0.036715 | TRUE |
| TXLNA | 1.461791997 | 4.313682714 | 0.700465954 | 2.086885148 | 0.036899 | TRUE |
| KCNH2 | 0.832604799 | 2.299300161 | 0.399160585 | 2.085889313 | 0.036989 | TRUE |
| ZDHHC23 | -1.30477819 | 0.27123269 | 0.62567834 | -2.085381747 | 0.037035 | TRUE |
| ARHGEF1 | 1.008994652 | 2.742842117 | 0.48384248 | 2.085378389 | 0.037035 | TRUE |
| HMGXB4 | -1.650068933 | 0.19203667 | 0.791516743 | -2.084692393 | 0.037097 | FALSE |
| PADI4 | 1.030222438 | 2.801688967 | 0.494213777 | 2.084568433 | 0.037108 | TRUE |
| ZNF382 | -1.106264455 | 0.330792346 | 0.531018581 | -2.083287655 | 0.037225 | TRUE |
| TARSL2 | -1.422455668 | 0.241121176 | 0.683045309 | -2.082520221 | 0.037295 | FALSE |

| | | | | | | |
|---|---|---|---|---|---|---|
| BAZ2B | -1.229474538 | 0.292446207 | 0.590383312 | -2.082502186 | 0.037297 | TRUE |
| JAK3 | 0.775841891 | 2.172420299 | 0.372609553 | 2.082184648 | 0.037326 | TRUE |
| WDR1 | 1.011398434 | 2.749443243 | 0.485775076 | 2.082030312 | 0.03734 | TRUE |
| IFFO1 | 1.048485681 | 2.853326996 | 0.50362841 | 2.081863653 | 0.037355 | TRUE |
| SNAP29 | 1.237165919 | 3.44583384 | 0.594301241 | 2.081715187 | 0.037368 | TRUE |
| SUPT3H | -1.128583278 | 0.323491229 | 0.5424434 | -2.080554909 | 0.037475 | TRUE |
| EDEM3 | -1.044211315 | 0.351969303 | 0.502132482 | -2.079553409 | 0.037567 | FALSE |
| CUZD1 | -1.168831945 | 0.310729679 | 0.562177522 | -2.079115403 | 0.037607 | TRUE |
| HLA-B | 1.115452504 | 3.050948433 | 0.536937397 | 2.077434931 | 0.037761 | TRUE |
| STT3B | -1.124118625 | 0.324938734 | 0.54145376 | -2.076111958 | 0.037884 | FALSE |
| MBTPS1 | 1.706676695 | 5.510617548 | 0.822095231 | 2.07600851 | 0.037893 | FALSE |
| CPOX | -1.307202965 | 0.270575809 | 0.629736689 | -2.075792926 | 0.037913 | TRUE |
| C15orf41 | -1.083537685 | 0.338396266 | 0.521992168 | -2.075773837 | 0.037915 | TRUE |
| GMFG | 1.261791191 | 3.53174185 | 0.607921084 | 2.075583862 | 0.037932 | TRUE |
| LAYN | -1.318297175 | 0.267590574 | 0.635333182 | -2.074969814 | 0.037989 | FALSE |
| FBXO22 | -1.287691278 | 0.275907041 | 0.620725602 | -2.074493584 | 0.038033 | FALSE |
| CD160 | -1.447137964 | 0.235242598 | 0.697644509 | -2.074320009 | 0.03805 | FALSE |
| TMEM11 | 1.21308147 | 3.363834252 | 0.584834476 | 2.074230435 | 0.038058 | FALSE |
| TMEM190 | 0.954452251 | 2.597247553 | 0.46027136 | 2.073672909 | 0.03811 | TRUE |
| TPM4 | 1.004898138 | 2.731629009 | 0.484605611 | 2.073641152 | 0.038113 | TRUE |
| USP14 | -1.290219946 | 0.275210245 | 0.622308618 | -2.073279895 | 0.038146 | TRUE |
| ZNF202 | -1.42140171 | 0.241375441 | 0.68575327 | -2.072759655 | 0.038195 | FALSE |
| USP6NL | -1.361373608 | 0.256308468 | 0.656803956 | -2.072724434 | 0.038198 | FALSE |
| ZSWIM3 | 1.452892634 | 4.275463999 | 0.70129003 | 2.071742891 | 0.038289 | TRUE |
| AP2A1 | 1.032420012 | 2.807852656 | 0.498726936 | 2.070110791 | 0.038442 | TRUE |
| PLCB2 | 0.979756885 | 2.663808553 | 0.473324424 | 2.069947875 | 0.038457 | TRUE |
| AUTS2 | -1.766617794 | 0.170910065 | 0.854001029 | -2.068636611 | 0.03858 | TRUE |
| PRKCG | 1.16710774 | 3.21268726 | 0.56419726 | 2.068616463 | 0.038582 | FALSE |
| SLC35F5 | -1.257274823 | 0.284428088 | 0.607854625 | -2.068380778 | 0.038604 | FALSE |
| ZNF292 | -1.164163172 | 0.312183797 | 0.563027201 | -2.067685485 | 0.03867 | FALSE |
| WASL | -1.063610702 | 0.345207117 | 0.514874395 | -2.065767327 | 0.03885 | FALSE |
| GMCL1 | -1.416633624 | 0.242529088 | 0.685909783 | -2.065335208 | 0.038891 | TRUE |
| EVA1A | -1.173260412 | 0.309356665 | 0.5682193 | -2.064802113 | 0.038942 | TRUE |
| RBM48 | -1.153360295 | 0.315574562 | 0.558597566 | -2.064742789 | 0.038947 | TRUE |
| ADAP2 | 1.51875871 | 4.566553257 | 0.73557966 | 2.064710039 | 0.03895 | FALSE |
| CHCHD3 | -1.411000849 | 0.243899055 | 0.683937711 | -2.063054611 | 0.039107 | TRUE |
| MEX3A | 1.02940087 | 2.799388135 | 0.499313179 | 2.061633686 | 0.039243 | TRUE |
| TCEB2 | 1.191054531 | 3.290549365 | 0.577755094 | 2.061521472 | 0.039253 | FALSE |
| C4orf45 | -1.357073732 | 0.257412935 | 0.658438475 | -2.061048653 | 0.039298 | TRUE |
| NDUFB2 | 1.371996056 | 3.943213719 | 0.665712396 | 2.060944131 | 0.039308 | TRUE |
| PTAFR | 1.206723584 | 3.342515222 | 0.585536364 | 2.060885809 | 0.039314 | TRUE |
| TRAPPC6B | -1.385757578 | 0.250134232 | 0.672700554 | -2.05999173 | 0.039399 | TRUE |
| PBX3 | -1.238592155 | 0.289791913 | 0.601315339 | -2.059804689 | 0.039417 | TRUE |
| DDX50 | -1.363334505 | 0.255806366 | 0.661914009 | -2.059685224 | 0.039429 | FALSE |

| | | | | | | |
|---|---|---|---|---|---|---|
| BRAF | -1.52883642 | 0.21678777 | 0.742497946 | -2.059044647 | 0.03949 | TRUE |
| ADH5 | -1.344843922 | 0.260580376 | 0.653307628 | -2.058515566 | 0.039541 | FALSE |
| HEATR6 | -1.331029496 | 0.264205123 | 0.646651602 | -2.058340987 | 0.039557 | TRUE |
| CTPS1 | -1.543692507 | 0.213590957 | 0.750061471 | -2.058087991 | 0.039582 | TRUE |
| RAC2 | 0.962438701 | 2.618073393 | 0.467786043 | 2.057433553 | 0.039645 | TRUE |
| NRL | 1.226042518 | 3.407716837 | 0.59623896 | 2.056293868 | 0.039754 | TRUE |
| MOB4 | 1.115396889 | 3.05077876 | 0.542515868 | 2.055970994 | 0.039785 | TRUE |
| IL36G | -1.206532369 | 0.299233113 | 0.586947374 | -2.055605704 | 0.039821 | TRUE |
| TAF1D | -1.15249949 | 0.315846327 | 0.560693652 | -2.055488741 | 0.039832 | FALSE |
| LTV1 | -1.39910855 | 0.246816891 | 0.680792979 | -2.055116008 | 0.039868 | TRUE |
| CUL2 | -1.494041942 | 0.224463551 | 0.72711603 | -2.054750384 | 0.039903 | FALSE |
| SP3 | -1.119248053 | 0.326525232 | 0.544717481 | -2.05473129 | 0.039905 | TRUE |
| PDHX | -1.222021248 | 0.294634036 | 0.595176288 | -2.053208895 | 0.040052 | TRUE |
| PTPRM | -0.711150967 | 0.491078657 | 0.346466927 | -2.052579657 | 0.040113 | TRUE |
| MRAP | 1.379902475 | 3.974513996 | 0.672322868 | 2.052440191 | 0.040127 | FALSE |
| TRAIP | 1.269691333 | 3.559753615 | 0.618785293 | 2.051909357 | 0.040178 | TRUE |
| ABR | 0.863868173 | 2.372319512 | 0.421011407 | 2.05188781 | 0.040181 | TRUE |
| CD72 | 1.584413874 | 4.876432338 | 0.772264407 | 2.051646898 | 0.040204 | TRUE |
| PRPF4B | -1.246928077 | 0.287386271 | 0.6080157 | -2.05081559 | 0.040285 | FALSE |
| NTRK1 | 1.097111831 | 2.995502003 | 0.53506042 | 2.050444752 | 0.040321 | TRUE |
| LINS1 | -1.207779847 | 0.298860058 | 0.589036855 | -2.050431712 | 0.040322 | TRUE |
| TECR | 0.963507709 | 2.62087363 | 0.469971036 | 2.050142743 | 0.040351 | FALSE |
| MMP11 | 0.90106478 | 2.462223441 | 0.43968051 | 2.049362571 | 0.040427 | TRUE |
| AUP1 | -1.16580823 | 0.311670658 | 0.568915572 | -2.049176165 | 0.040445 | FALSE |
| ATPAF2 | 1.447646121 | 4.25309146 | 0.70652612 | 2.048963344 | 0.040466 | TRUE |
| FAM129B | 1.075267724 | 2.930777434 | 0.525015462 | 2.048068678 | 0.040553 | TRUE |
| GUCA2A | 1.111935296 | 3.040236462 | 0.543212867 | 2.046960527 | 0.040662 | FALSE |
| C17orf89 | 1.208734593 | 3.349243814 | 0.590543901 | 2.04681581 | 0.040676 | TRUE |
| POMT2 | -1.679701242 | 0.186429665 | 0.820962454 | -2.046014691 | 0.040755 | FALSE |
| TXNDC16 | -1.231449953 | 0.291869074 | 0.602172762 | -2.045011052 | 0.040854 | FALSE |
| STXBP4 | -1.116907894 | 0.327290248 | 0.546199327 | -2.044872337 | 0.040867 | FALSE |
| NUP54 | -1.214660949 | 0.296810631 | 0.594067965 | -2.044649806 | 0.040889 | FALSE |
| NR1D1 | 1.02530598 | 2.787948385 | 0.501462007 | 2.04463342 | 0.040891 | FALSE |
| CCDC153 | 1.129545385 | 3.094249489 | 0.552592294 | 2.044084577 | 0.040945 | FALSE |
| MYADML2 | 1.239101864 | 3.452511249 | 0.60621333 | 2.044002997 | 0.040953 | TRUE |
| HAP1 | 1.227980476 | 3.414327252 | 0.601045626 | 2.043073643 | 0.041045 | FALSE |
| GPR135 | -1.117873455 | 0.326974382 | 0.547222554 | -2.042813195 | 0.041071 | FALSE |
| ARL4D | 1.173191276 | 3.23229133 | 0.574399449 | 2.04246588 | 0.041105 | FALSE |
| PRR5 | 1.075705903 | 2.932061921 | 0.526770216 | 2.042078066 | 0.041144 | FALSE |
| UPK3A | 0.912648811 | 2.490911757 | 0.44692496 | 2.042062747 | 0.041145 | TRUE |
| RIOK2 | -1.138572832 | 0.320275783 | 0.557609306 | -2.041882767 | 0.041163 | TRUE |
| ZNF614 | -1.193229833 | 0.303240265 | 0.584617717 | -2.041042887 | 0.041247 | TRUE |
| SETD9 | -1.326202469 | 0.265483531 | 0.649821604 | -2.040871617 | 0.041264 | FALSE |
| CES2 | 1.222196892 | 3.394637198 | 0.598950512 | 2.040564067 | 0.041294 | FALSE |

| | | | | | | |
|---|---|---|---|---|---|---|
| SCP2 | -1.126198842 | 0.324263494 | 0.551974677 | -2.040308892 | 0.04132 | TRUE |
| DNAJC1 | -1.438407263 | 0.237305423 | 0.705813486 | -2.037942447 | 0.041556 | TRUE |
| RC3H2 | -1.477130585 | 0.228291814 | 0.725145227 | -2.037013454 | 0.041649 | FALSE |
| KIR3DL3 | 1.122079876 | 3.071235354 | 0.550933186 | 2.036689576 | 0.041681 | FALSE |
| UQCRB | -1.142892379 | 0.318895321 | 0.561206975 | -2.036489975 | 0.041701 | FALSE |
| CORO6 | 1.108465499 | 3.029705739 | 0.54454397 | 2.03558493 | 0.041792 | TRUE |
| DAAM2 | -1.092989545 | 0.33521286 | 0.537100714 | -2.034980624 | 0.041853 | TRUE |
| FHOD1 | 0.840435601 | 2.317376208 | 0.413204759 | 2.033944633 | 0.041957 | TRUE |
| DHRS3 | 1.395073804 | 4.035272381 | 0.686315346 | 2.032700876 | 0.042083 | FALSE |
| AFG3L2 | -1.528186715 | 0.216928664 | 0.75204289 | -2.032047287 | 0.042149 | FALSE |
| RPUSD4 | -1.128702399 | 0.323452697 | 0.555516957 | -2.031805482 | 0.042173 | FALSE |
| PDE7B | -1.251067401 | 0.286199144 | 0.61579872 | -2.031617412 | 0.042192 | TRUE |
| SULT1A2 | 1.180313802 | 3.255395592 | 0.581152083 | 2.030989541 | 0.042256 | TRUE |
| TBCEL | -1.143177914 | 0.318804278 | 0.563059119 | -2.030298196 | 0.042326 | TRUE |
| HFM1 | -0.931223482 | 0.394071276 | 0.458682199 | -2.030215006 | 0.042335 | TRUE |
| SOBP | -1.45503973 | 0.233391091 | 0.716810383 | -2.029880935 | 0.042369 | TRUE |
| CCDC94 | 1.121223946 | 3.068607715 | 0.552437806 | 2.029593076 | 0.042398 | TRUE |
| TSPAN16 | 1.076939905 | 2.935682325 | 0.530742571 | 2.029119131 | 0.042446 | FALSE |
| KDM3B | 1.854140739 | 6.386208476 | 0.913834792 | 2.028967113 | 0.042462 | TRUE |
| MS4A12 | 1.244567483 | 3.471433022 | 0.6136589 | 2.028109563 | 0.042549 | TRUE |
| AC006372.1 | -1.014428949 | 0.362609439 | 0.500330562 | -2.027517459 | 0.04261 | TRUE |
| SAAL1 | -1.298029046 | 0.27306947 | 0.640375211 | -2.026982031 | 0.042664 | TRUE |
| HEATR4 | -1.226196569 | 0.293406409 | 0.604979441 | -2.026840063 | 0.042679 | TRUE |
| GZMA | 1.120472127 | 3.066301545 | 0.552958064 | 2.026323876 | 0.042732 | FALSE |
| ACTN1 | 1.233169259 | 3.432089498 | 0.608814932 | 2.025524003 | 0.042814 | TRUE |
| PIGF | -1.094727952 | 0.33463063 | 0.540484325 | -2.025457357 | 0.04282 | FALSE |
| ZSCAN10 | 1.070893382 | 2.91798521 | 0.528869458 | 2.024872801 | 0.04288 | TRUE |
| NUP35 | -1.258469685 | 0.284088439 | 0.621580755 | -2.024627813 | 0.042906 | FALSE |
| TCAP | 0.919273178 | 2.507467245 | 0.454313362 | 2.023434163 | 0.043028 | TRUE |
| TBX4 | 1.098820931 | 3.000625991 | 0.543049953 | 2.023425148 | 0.043029 | TRUE |
| SLC1A4 | 1.424437313 | 4.155518927 | 0.70430202 | 2.0224808 | 0.043127 | TRUE |
| CLPP | 1.199813379 | 3.319497377 | 0.593475339 | 2.021673522 | 0.04321 | FALSE |
| UCKL1 | 0.704371869 | 2.022575844 | 0.34841714 | 2.021633806 | 0.043214 | TRUE |
| RAB25 | 1.189240052 | 3.284584147 | 0.588363354 | 2.021268055 | 0.043252 | FALSE |
| COA1 | -1.568270726 | 0.208405261 | 0.775909658 | -2.02120274 | 0.043259 | TRUE |
| CYBA | 0.667982154 | 1.950297947 | 0.330743216 | 2.019639775 | 0.043421 | FALSE |
| RAB22A | -1.356503728 | 0.257559703 | 0.671686516 | -2.019548846 | 0.04343 | TRUE |
| TSKS | 1.288785631 | 3.628377689 | 0.638382238 | 2.018830653 | 0.043505 | TRUE |
| MZF1 | 1.0245942 | 2.785964686 | 0.507624661 | 2.01840903 | 0.043549 | FALSE |
| ATL2 | -1.44422078 | 0.235929846 | 0.715605969 | -2.018178778 | 0.043573 | FALSE |
| BCL2L2-PABPN1 | 1.244885169 | 3.472536023 | 0.616837409 | 2.018173917 | 0.043573 | TRUE |
| MRGPRD | 1.052105273 | 2.863673591 | 0.521550139 | 2.017265829 | 0.043668 | FALSE |
| PTPN21 | -1.294786047 | 0.273956472 | 0.641940875 | -2.016986451 | 0.043697 | TRUE |
| STEAP1B | -1.105778876 | 0.330953011 | 0.548301308 | -2.016735796 | 0.043723 | TRUE |

| | | | | | | |
|---|---|---|---|---|---|---|
| PALMD | -0.976024754 | 0.376806022 | 0.48405392 | -2.016355437 | 0.043763 | TRUE |
| INSIG1 | 1.308525682 | 3.700713658 | 0.649095649 | 2.015921205 | 0.043808 | TRUE |
| BEND7 | -1.492016025 | 0.224918756 | 0.74012256 | -2.015903994 | 0.04381 | FALSE |
| PVALB | 0.976029468 | 2.653897908 | 0.484305573 | 2.015317441 | 0.043871 | TRUE |
| U2SURP | -1.394209936 | 0.248028918 | 0.691867311 | -2.01514064 | 0.04389 | TRUE |
| SLC25A36 | -1.077646471 | 0.340395715 | 0.535079078 | -2.013994782 | 0.04401 | FALSE |
| TUBA1C | 1.555382363 | 4.736897395 | 0.772534919 | 2.013348945 | 0.044078 | TRUE |
| TBCK | -1.112865917 | 0.328615825 | 0.552955126 | -2.012579078 | 0.044159 | TRUE |
| PTGDS | 0.710057051 | 2.034107303 | 0.352845557 | 2.012373505 | 0.044181 | TRUE |
| PECR | -1.301064809 | 0.272241753 | 0.646637763 | -2.012045822 | 0.044215 | TRUE |
| TDG | -1.257186804 | 0.284453124 | 0.624968988 | -2.011598699 | 0.044262 | FALSE |
| LAPTM5 | 1.212973674 | 3.363471664 | 0.603003796 | 2.011552303 | 0.044267 | TRUE |
| UBALD2 | 1.114665081 | 3.048546991 | 0.554171799 | 2.01140708 | 0.044282 | FALSE |
| CA9 | 1.161829581 | 3.195774861 | 0.577740265 | 2.010989456 | 0.044327 | TRUE |
| B3GNT9 | 0.679467137 | 1.972826206 | 0.337920625 | 2.010729995 | 0.044354 | FALSE |
| NEK6 | 1.050542424 | 2.859201596 | 0.522579168 | 2.010302911 | 0.044399 | TRUE |
| ERLIN1 | -1.271421224 | 0.280432781 | 0.632921645 | -2.008812992 | 0.044557 | FALSE |
| BROX | 1.205652228 | 3.338936114 | 0.60019324 | 2.008773421 | 0.044561 | TRUE |
| LRWD1 | 0.762846688 | 2.144371898 | 0.37986471 | 2.008206259 | 0.044621 | TRUE |
| CALML5 | 0.794912788 | 2.21424788 | 0.395915616 | 2.007783368 | 0.044666 | TRUE |
| TMEM259 | 0.574207377 | 1.77572249 | 0.286096997 | 2.007037415 | 0.044746 | TRUE |
| PRB1 | -0.893838859 | 0.409082325 | 0.44538444 | -2.006892875 | 0.044761 | FALSE |
| COBL | -1.191868509 | 0.303653355 | 0.593930833 | -2.006746312 | 0.044777 | TRUE |
| ANO8 | 0.728216192 | 2.071382362 | 0.363009921 | 2.006050388 | 0.044851 | TRUE |
| FIGN | -0.412116736 | 0.662246963 | 0.205471914 | -2.005708362 | 0.044887 | TRUE |
| DDX10 | -1.18531451 | 0.305650034 | 0.591064682 | -2.005388829 | 0.044921 | TRUE |
| MARCH7 | -1.191103978 | 0.303885596 | 0.594370234 | -2.003976494 | 0.045073 | FALSE |
| ZNF784 | 0.87136896 | 2.390180676 | 0.434870187 | 2.003744994 | 0.045097 | FALSE |
| PLPP1 | -1.307982207 | 0.270365047 | 0.65281756 | -2.003595317 | 0.045113 | TRUE |
| DCDC2 | -0.98705043 | 0.3726743 | 0.492807392 | -2.002913199 | 0.045187 | TRUE |
| FLOT1 | 1.256306782 | 3.512425349 | 0.627457848 | 2.002217019 | 0.045261 | FALSE |
| LSM4 | 0.853538221 | 2.347939702 | 0.426451561 | 2.001489266 | 0.04534 | TRUE |
| FBXO5 | -1.109370697 | 0.329766419 | 0.554321337 | -2.001313357 | 0.045359 | FALSE |
| CFAP36 | 1.401594987 | 4.061673118 | 0.700424564 | 2.001064868 | 0.045385 | TRUE |
| SAMD5 | -0.767913275 | 0.463980258 | 0.384036562 | -1.999583768 | 0.045545 | TRUE |
| RHOU | -1.326714067 | 0.265347745 | 0.663730199 | -1.99887555 | 0.045622 | FALSE |
| MYOM2 | -0.945386232 | 0.388529479 | 0.473241141 | -1.997683953 | 0.045751 | TRUE |
| PHC3 | -1.357558399 | 0.257288206 | 0.67959374 | -1.997602859 | 0.04576 | TRUE |
| SSTR2 | -1.246836434 | 0.287412609 | 0.624615997 | -1.996164748 | 0.045916 | TRUE |
| ELMOD2 | -1.13705703 | 0.320761626 | 0.569833042 | -1.995421371 | 0.045997 | TRUE |
| GAS2 | -0.873953724 | 0.417298401 | 0.438014893 | -1.995260294 | 0.046014 | TRUE |
| RPP30 | -1.200402496 | 0.301073007 | 0.601974469 | -1.994108652 | 0.04614 | TRUE |
| LTBP4 | 0.955005779 | 2.598685601 | 0.479041992 | 1.993574249 | 0.046199 | FALSE |
| NDUFS8 | 0.922109331 | 2.514588901 | 0.462563997 | 1.993474064 | 0.04621 | TRUE |

| | | | | | | |
|---|---|---|---|---|---|---|
| MLLT4 | -0.774704637 | 0.460839876 | 0.388660767 | -1.99326689 | 0.046232 | TRUE |
| AICDA | 1.295110032 | 3.651397721 | 0.64992 | 1.99272223 | 0.046292 | TRUE |
| NT5DC1 | -1.407958289 | 0.244642262 | 0.706900478 | -1.991734809 | 0.0464 | FALSE |
| GPX5 | 1.216274873 | 3.374593501 | 0.610700819 | 1.991605113 | 0.046414 | FALSE |
| OSBPL9 | -1.558832761 | 0.210381494 | 0.782810254 | -1.991329001 | 0.046445 | FALSE |
| SLC27A3 | 1.103684328 | 3.015254772 | 0.554319814 | 1.991060576 | 0.046474 | TRUE |
| CHN1 | -1.326547873 | 0.265391848 | 0.666271883 | -1.991000831 | 0.046481 | TRUE |
| BIRC7 | 0.826130041 | 2.284460841 | 0.415037939 | 1.990492826 | 0.046537 | FALSE |
| TPD52 | -1.472374716 | 0.229380125 | 0.73993006 | -1.989883632 | 0.046604 | FALSE |
| LETM2 | -1.253022911 | 0.285640026 | 0.629890359 | -1.989271455 | 0.046671 | FALSE |
| PPM1D | -1.55515784 | 0.211156051 | 0.781786923 | -1.989234911 | 0.046675 | TRUE |
| NLRP3 | 1.211332049 | 3.357954635 | 0.609032726 | 1.988944104 | 0.046707 | TRUE |
| DEF8 | 0.876606164 | 2.402731376 | 0.440776031 | 1.988779113 | 0.046726 | FALSE |
| ANKRD65 | 0.746120137 | 2.10880226 | 0.375576414 | 1.98660009 | 0.046967 | FALSE |
| KIAA1109 | -0.992503367 | 0.370647662 | 0.499625102 | -1.986496201 | 0.046978 | TRUE |
| SREK1IP1 | -1.112783832 | 0.3286428 | 0.560211401 | -1.986364129 | 0.046993 | TRUE |
| CXCL6 | -1.063786078 | 0.345146582 | 0.535725772 | -1.985691437 | 0.047068 | TRUE |
| CDK9 | 1.027589645 | 2.794322401 | 0.517533067 | 1.985553601 | 0.047083 | TRUE |
| SYNGR1 | 1.084807385 | 2.958869842 | 0.546483704 | 1.985068132 | 0.047137 | TRUE |
| KLHL2 | -1.107924644 | 0.330243623 | 0.558231587 | -1.984704324 | 0.047177 | FALSE |
| C2orf40 | -1.268397198 | 0.2812821 | 0.639107242 | -1.984639062 | 0.047185 | FALSE |
| RASAL1 | 1.17093647 | 3.22501135 | 0.590073627 | 1.98439045 | 0.047212 | TRUE |
| PMPCB | -1.1412853 | 0.319408223 | 0.575506707 | -1.983096435 | 0.047357 | FALSE |
| TMUB2 | 1.131025424 | 3.098832488 | 0.570599075 | 1.982171851 | 0.04746 | FALSE |
| AP003733.1 | 1.196980561 | 3.310107151 | 0.604624082 | 1.979710361 | 0.047736 | TRUE |
| OTUD6B | -1.041314666 | 0.352990312 | 0.526059807 | -1.979460608 | 0.047764 | FALSE |
| SLC33A1 | -1.25473218 | 0.285152207 | 0.633901666 | -1.979379842 | 0.047773 | FALSE |
| BCL3 | 0.940420852 | 2.561059018 | 0.475271064 | 1.978704201 | 0.047849 | TRUE |
| DESI1 | -1.323451264 | 0.266214937 | 0.668880573 | -1.978606224 | 0.04786 | FALSE |
| GGH | -1.067066025 | 0.344016374 | 0.539372293 | -1.978347867 | 0.047889 | FALSE |
| POLA2 | 1.364099661 | 3.912199159 | 0.689561871 | 1.978212136 | 0.047905 | TRUE |
| KLK6 | 1.093805433 | 2.985614037 | 0.553248782 | 1.97705891 | 0.048035 | TRUE |
| CLDN3 | 0.606900032 | 1.834734954 | 0.307143137 | 1.975951794 | 0.04816 | FALSE |
| NKAIN2 | -0.710333522 | 0.491480251 | 0.359927902 | -1.97354392 | 0.048434 | TRUE |
| LIM2 | -1.268895509 | 0.281141969 | 0.643098398 | -1.973096984 | 0.048485 | FALSE |
| ATP5E | -1.114537505 | 0.328066973 | 0.565322251 | -1.971508293 | 0.048666 | FALSE |
| PLEKHB1 | 1.176400652 | 3.242681632 | 0.596870043 | 1.970949396 | 0.04873 | FALSE |
| GREB1L | -1.113940346 | 0.32826294 | 0.565386792 | -1.97022704 | 0.048812 | TRUE |
| MET | -0.946144448 | 0.388235002 | 0.480259616 | -1.970068723 | 0.04883 | TRUE |
| CCDC14 | -0.938293751 | 0.391294913 | 0.476371568 | -1.969667825 | 0.048876 | FALSE |
| AZU1 | 0.87925768 | 2.409110712 | 0.446552841 | 1.96898911 | 0.048954 | TRUE |
| INVS | -1.28109924 | 0.277731839 | 0.650653641 | -1.968941938 | 0.04896 | TRUE |
| LILRA5 | 0.926418529 | 2.525448141 | 0.470534577 | 1.968863871 | 0.048969 | FALSE |
| MERTK | -1.752038302 | 0.1734201 | 0.890186435 | -1.968170074 | 0.049048 | FALSE |

| | | | | | | |
|---|---|---|---|---|---|---|
| PLEKHH1 | -1.376496234 | 0.252461572 | 0.700090363 | -1.966169377 | 0.049279 | FALSE |
| KITLG | -1.029772086 | 0.357088337 | 0.523945981 | -1.965416521 | 0.049366 | TRUE |
| ZNF70 | 1.041381895 | 2.833129399 | 0.529870921 | 1.965350153 | 0.049374 | TRUE |
| ZEB1 | -0.878654725 | 0.415341284 | 0.447123071 | -1.965129475 | 0.049399 | TRUE |
| C5AR1 | 1.195505642 | 3.305228609 | 0.608590752 | 1.964383517 | 0.049486 | TRUE |
| RP1-66C13.4 | -1.468245831 | 0.230329167 | 0.747654503 | -1.963802565 | 0.049553 | FALSE |
| HIVEP1 | -1.369289528 | 0.25428756 | 0.697319077 | -1.963648456 | 0.049571 | TRUE |
| G6PC3 | -1.129247197 | 0.323276528 | 0.575087874 | -1.963608082 | 0.049576 | TRUE |
| UBXN7 | -1.760203398 | 0.172009874 | 0.896545077 | -1.963318346 | 0.049609 | FALSE |
| MAPKBP1 | 1.596379795 | 4.935133849 | 0.813326787 | 1.96277784 | 0.049672 | TRUE |
| PSMD1 | -1.177522456 | 0.308040979 | 0.600320124 | -1.961490891 | 0.049822 | TRUE |
| PIDD1 | 0.851316031 | 2.342727928 | 0.434050152 | 1.961331028 | 0.04984 | TRUE |
| ME2 | -1.413406753 | 0.243312963 | 0.720722503 | -1.961097019 | 0.049868 | TRUE |
| APBB2 | -0.887180636 | 0.411815174 | 0.452489614 | -1.960665191 | 0.049918 | TRUE |
| SKOR1 | 0.777625495 | 2.176298493 | 0.39665844 | 1.960441063 | 0.049944 | FALSE |
| CTDSP2 | 1.363545092 | 3.910030178 | 0.695557996 | 1.960361466 | 0.049954 | TRUE |
| PARG | -1.207457663 | 0.298956362 | 0.615950185 | -1.960317071 | 0.049959 | FALSE |
